# *NPRL3*: Direct Effects on Human Phenotypic Variability, mTOR Signaling, Subcellular mTOR Localization, Cortical Lamination, and Seizure Susceptibility

**DOI:** 10.1101/2020.12.11.421214

**Authors:** Philip H. Iffland, Mariah E. Everett, Katherine M. Cobb-Pitstick, Lauren E. Bowser, Allan E. Barnes, Janice K. Babus, Andrea Romanowski, Marianna Baybis, Erik G. Puffenberger, Claudia Gonzaga-Jauregui, Alexandros Poulopoulos, Vincent J. Carson, Peter B. Crino

## Abstract

Nitrogen Permease Regulator Like 3 *(NPRL3)* variants are associated with malformations of cortical development (MCD) and epilepsy. We report a large (n=133) founder *NPRL3* (c.349delG, p.Glu117LysFS) pedigree dating to 1727, with heterogeneous epilepsy and MCD phenotypes. Whole exome analysis in individuals with and without seizures in this cohort did not identify a genetic modifier to explain the variability in seizure phenotype. Then as a strategy to investigate the developmental effects of NPRL3 loss in human brain, we show that CRISPR/Cas9 *Nprl3* knockout (KO) in Neuro2a cells (N2aC) *in vitro* causes mechanistic target of rapamycin (mTOR) pathway hyperactivation, cell soma enlargement, and excessive cellular aggregation. Amino acid starvation caused mTOR inhibition and cytoplasmic mTOR localization in wildtype cells, whereas following *Nprl3* KO, mTOR remained inappropriately localized on the lysosome and activated, evidenced by persistent ribosomal S6 and 4E-BP1 phosphorylation, demonstrating that *Nprl3* loss decouples mTOR activation from metabolic state. *Nprl3* KO by *in utero* electroporation in fetal (E14) mouse cortex resulted in mTOR-dependent cortical dyslamination with ectopic neurons in subcortical white matter. EEG recordings of these mice showed hyperexcitability in the electroporated hemisphere. *NPRL3* variants are linked to a highly variable clinical phenotype likely as a consequence of mTOR-dependent effects on cell structure, cortical development, and network organization.

## Introduction

Malformations of cortical development (MCD) are common causes of medically refractory epilepsy. Germline and somatic variants in mTOR pathway genes (MPG) are the most common cause of MCD^1^. The mTOR pathway plays a pivotal role in cerebral cortical development, coordinating proliferation, size, polarization, and migration in fetal cortex^2^ and MPG variants leading to constitutive mTOR activation (‘mTORopathies’) provide the presumed mechanism for several MCD subtypes. Focal cortical dysplasia II (FCDII) is an MCD highly associated with intractable seizures defined by disorganized cerebral cortical lamination, enlarged (cytomegalic) dysmorphic neurons, and heterotopic neurons in the white matter^3^. Variants in genes encoding components of the GTPase Activating Protein (GAP) Activity Towards Rags Complex 1 (GATOR1), including DEP domain containing 5 (*DEPDC5*), Nitrogen Permease Regulator Like 2 (*NPRL2*), and Nitrogen Permease Regulator Like 3 (*NPRL3*)*^4–6^,* are the most common MPG variants associated with FCDII^7^ and hemimegalencephaly (HME), a MCD defined by a hemispheric enlargement and FCD-type II histopathology ^8,9^.

In non-neural cell types, the GATOR1 complex modulates metabolic mTOR pathway activity in response to intracellular amino acid levels by governing translocation of mTORC1 (mTOR complex 1, mTOR bound to raptor) to and from the lysosomal membrane ^10,11^. In replete amino acid conditions, GATOR1 is inhibited, releasing mTORC1 to physically interact with its binding partners e.g., Rheb, on the lysosomal surface in active conformation. However, when intracellular amino acids are low, GATOR1 alters the nucleotide bound state of the Rag proteins and prevents translocation of mTORC1 to the lysosomal surface (remaining cytoplasmic) thereby inhibiting mTOR activity^12^. GATOR1 subunits are highly expressed in brain ^13,14^ and the effects of for example, *DEPDC5* variants on cell size, motility, and firing as well as cortical lamination have been shown in several experimental models^15–20^ and in human brain specimens ^4,7^. While human tissue studies show a link between *NPRL3* variants and mTOR activation^9,21–23^, surprisingly few studies have investigated the role of *NPRL3* in neuronal morphology, cortical lamination, and cortical circuitry.

We report the largest and genealogically oldest known *NPRL3* patient pedigree and show that despite a common founder variant, clinical findings among affected individuals were highly heterogeneous with a spectrum ranging from normal brain imaging to HME and seizure freedom to intractable epilepsy^21^. As a strategy to define mechanisms for clinical phenotypic heterogeneity, whole exome analysis was performed in this cohort but did not identify a gene modifier for seizure phenotype. To assess the role of Nprl3 in murine fetal brain development, CRISPR/Cas9 KO of *Nprl3 in vitro* resulted in mTOR signaling hyperactivation and mTOR-dependent abnormalities in cell size and subcellular translocation of mTOR to the lysosomal membrane. Focal *Nprl3* KO using *in utero* electroporation (IUE) in fetal mice caused mTOR-dependent cortical lamination defects and increased seizure susceptibility in the post-natal period.

## Results

### NPRL3 Pedigree

Old Order Mennonite probands (n=133) heterozygous for the *NPRL3* c.349delG, p.Glu117LysFS variant^22^ and born between 1934 and 2020 were assembled across eight states: Pennsylvania (n=83), Ohio (n=25), New York (n=8), Wisconsin (n=7), Missouri (n=6), Tennessee (n=2), Indiana (n=1), and Kentucky (n=1). All patients were connected through a twelve-generation pedigree to a founder mutation likely originating in an Old Order Mennonite couple from Pennsylvania born in 1727 and 1728 (Fig. 1). Analysis of our Mennonite exome control database (Clinic for Special Children, Strasburg, PA) revealed that the *NPRL3* variant had a minor allele frequency of 0.725% (corresponding carrier frequency of 1.44%) in the Weaverland and Groffdale Conference Mennonite population.

**Figure 1:**
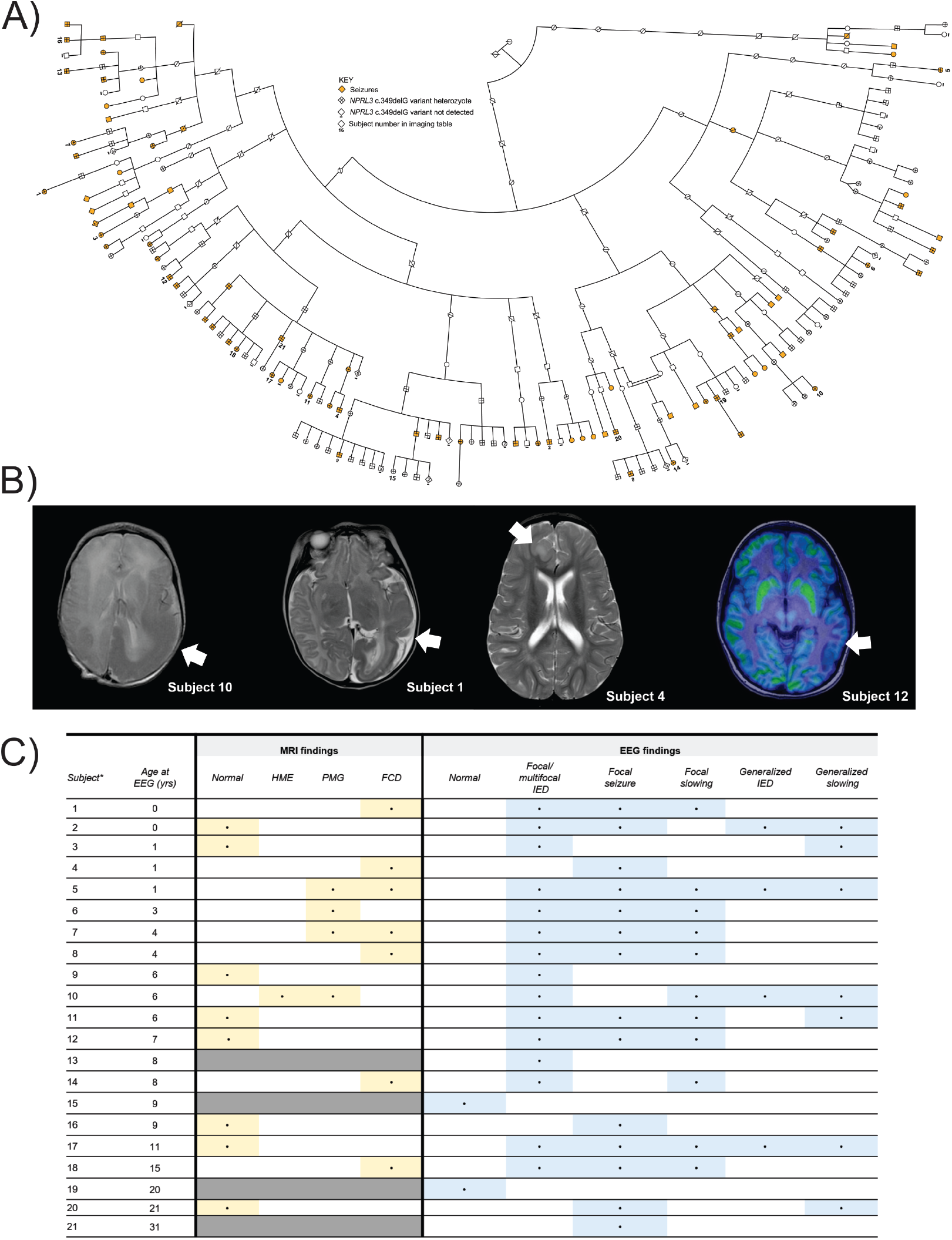
*NPRL3* founder mutation segregates in a large Old Order Mennonite pedigree. (A) One hundred and thirty-three Mennonite *NPRL3* c.349delG variant heterozygotes were traced to a common founder couple (born 1727 and 1728) through 12 generations. Heterozygotes are denoted with a cross-mark, while patients in which the variant was not detected are denoted with a (-) below their symbol. Patients with seizures are denoted in orange. A number below a patient symbol corresponds to the subject number listed in (C) for EEG and MRI results. Of note, only one patient who did not have the *NPRL3* variant reported seizures. *Number inside of diamond represents condensed sibling numbers to conserve pedigree space.* (B) MRI and PET /MRI images for subjects 10, 1, 4, and 12. Subject 10 (*far left)* at age 3 days, T2-weighted image showing left hemimegalencephaly with associated cortical thickening. Subject 1 (*second from left)* at age 7 weeks, T2-weighted image showing large left hemispheric cortical dysplasia involving the parietal, temporal, and occipital lobes. Subject 4 (*second from right)* at age 1 year 7 months, T2-weighted image showing focal cortical dysplasia in the right frontal lobe. Subject 12 (*far right)* at age 7 years 11 months, PET/MRI showing large area of hypometabolism in the left posterior temporal cortex in a patient without obvious focal cortical dysplasia on previous imaging. (C) MRI and EEG data were available for 17 patients. One patient (subject 10) had hemimegalencephaly (HME) on MRI, while seven patients had focal cortical dysplasia (2 with polymicrogyria as well). One patient had polymicrogyria (PMG) only and eight individuals had normal MRIs despite a clinical and/or electrographic diagnosis of seizures. On EEG, patients showed both generalized and focal abnormalities. Of the two patients with normal EEGs, subject 15 underwent EEG as part of clinical care and subject 19 had a clinical diagnosis of seizures.

Among the cohort of 133 *NPRL3* mutation heterozygotes, 48 (36.1%) had a history of seizures while 85 had no seizure history. Of the 48 *NPRL3* individuals with seizures, 42 completed structured medical interviews (6 were excluded from interviews due to a dual diagnosis). Affected individuals showed markedly heterogeneous epilepsy phenotypes. The median age of seizure onset was 5 years but ranged from one day to 37 years. The median peak seizure frequency was 4.5 seizures per day but ranged from 4 per year to 100 per day. Six individuals had a history of infantile spasms and 1 reported exclusively febrile seizures. Six patients underwent epilepsy surgery. The median number of concurrent anti-seizure medications was 2 and ranged from 0 to 4. While *DEPDC5* variants have been associated with sudden unexplained death in epilepsy (SUDEP), there were no reported cases of possible or confirmed SUDEP within our *NPRL3* pedigree.

Within the cohort, the estimated penetrance of seizures was 28% (penetrance may be an underestimate as some patients may yet develop seizures later in life). To reduce the likelihood of overestimating the penetrance, out of the cohort of 133 subjects, we identified 10 families in which an entire sibship was tested for the *NPRL3* mutation, thus all *NPRL3* heterozygotes without seizures were identified. In these sibships, there were 39 heterozygotes, 11 of whom had seizures.

MRI data were available for 17 *NPRL3* mutation heterozygotes and EEG data for 21, both showing highly heterogeneous results (Fig. 1). MRI findings included normal brain structure in 8 individuals and MCD (polymicrogyria in 3, FCD in 7, and HME in 1) in 11 (Fig. 1). Two of 21 individuals had normal EEGs (these had no MRI available). EEGs showed both focal (spikes, sharp waves) and generalized (generalized spike and wave discharges and slowing) abnormalities. The most common pattern observed was focal or multi-focal inter-ictal discharges in 15 patients across a range of MRI findings, including normal brain imaging. One individual in the pedigree with a single clinical seizure did not harbor the *NPRL3* c.349delG, p.Glu117LysFS variant. This *NPRL3* variant was detected in her mother and 1 sister, both with no clinical phenotype, and another sister (subject 17) with epilepsy, a normal MRI, and abnormal EEG. A third sister was phenotypically normal and had negative *NPRL3* testing.

#### Gene Modifier Analysis

In view of the clinical heterogeneity of the pedigree, we performed whole exome sequencing through the Regeneron Genetics Center on 83 *NPRL3* heterozygotes to investigate the possibility of genetic modifiers affecting *NPRL3* c.349delG penetrance. Genome-wide association analysis of the *NPRL3* c.349delG heterozygotes against exome data from 835 Old Order Mennonite control samples was completed. For each variant identified in the *NPRL3* exome cohort, we calculated a chi-square statistic to assess allele frequency differences between patients and controls. The results (Supp. Table A) demonstrated that the *NPRL3* c.349delG variant exhibited the greatest statistical allele frequency deviation out of 230,656 variants (as expected). The top 25 also contained 9 additional variants from distal chromosome 16p13 that demonstrated highly significant chi-square values. This, however, was not surprising as these variants are in linkage disequilibrium with the *NPRL3* c.349delG variant, and they are frequently co-inherited on the same disease haplotype.

Next, we split our cohort into 3 partially-overlapping clinical groups: “no seizures” (n=37), “seizures” (monthly seizures; n=24), and “severe seizures” (weekly seizures; n=10). We performed two separate whole genome-wide analyses comparing the “no seizure” group to both the “seizure” group (Supp. Table B) and the “severe seizure” group (Supp. Table C). The tables list all variants that passed filtering (n=71,199) and demonstrated significant chi-square values (>3.84). Using both the chi-square statistic and the genotype classes, we failed to find a single common variant that could explain the differences in epilepsy penetrance and severity in our *NPRL3* cohort. Of course, while it is possible that multiple polygenic modifiers may act independently in each individual or sibship that our sample size is likely too small to detect, these data are consistent with an alternate monogenic mechanism through which heterozygous *NPRL3* genotypes result in the partially penetrant and clinically heterogeneous phenotype.

### mTOR Pathway Activation

Enhanced phosphorylation of ribosomal S6 protein at serine 240/244 (PS6; 240/244) is a well-defined effect of mTOR pathway hyperactivation as PS6 is a downstream target of mTOR. Elevated PS6 has been reported in human *NPRL3* variant brain tissue specimens by immunohistochemistry^15,23^ and in experimental models of *Nprl3* loss^15,24^. Two gRNAs (*Nprl3* (A) and *Nprl3* (B); -GAGGTGTCTGCTATGGCTGA- and -AATTGCTACTGTCCTGCAGC-, respectively), targeting exon 5 of *Nprl3* were transfected into N2aC lines to achieve *Nprl3* knockout (KO) (Fig. 2A). Increased PS6 levels were detected in lysates from *Nprl3 (A)* and *(B)* KO cell lines compared to scramble sequence transfected and WT cell lines grown in complete media (Fig. 2B; densitometry data in Supp. Fig. 1). PS6 levels were reduced to WT levels in *Nprl3* KO lines treated with rapamycin (150 nM) or abolished with the ATP-competitive mTOR inhibitor torin1 (100 nM) for 60 minutes in complete media (Fig. 2B; densitometry data in Supp. Fig. 1).

**Figure 2:**
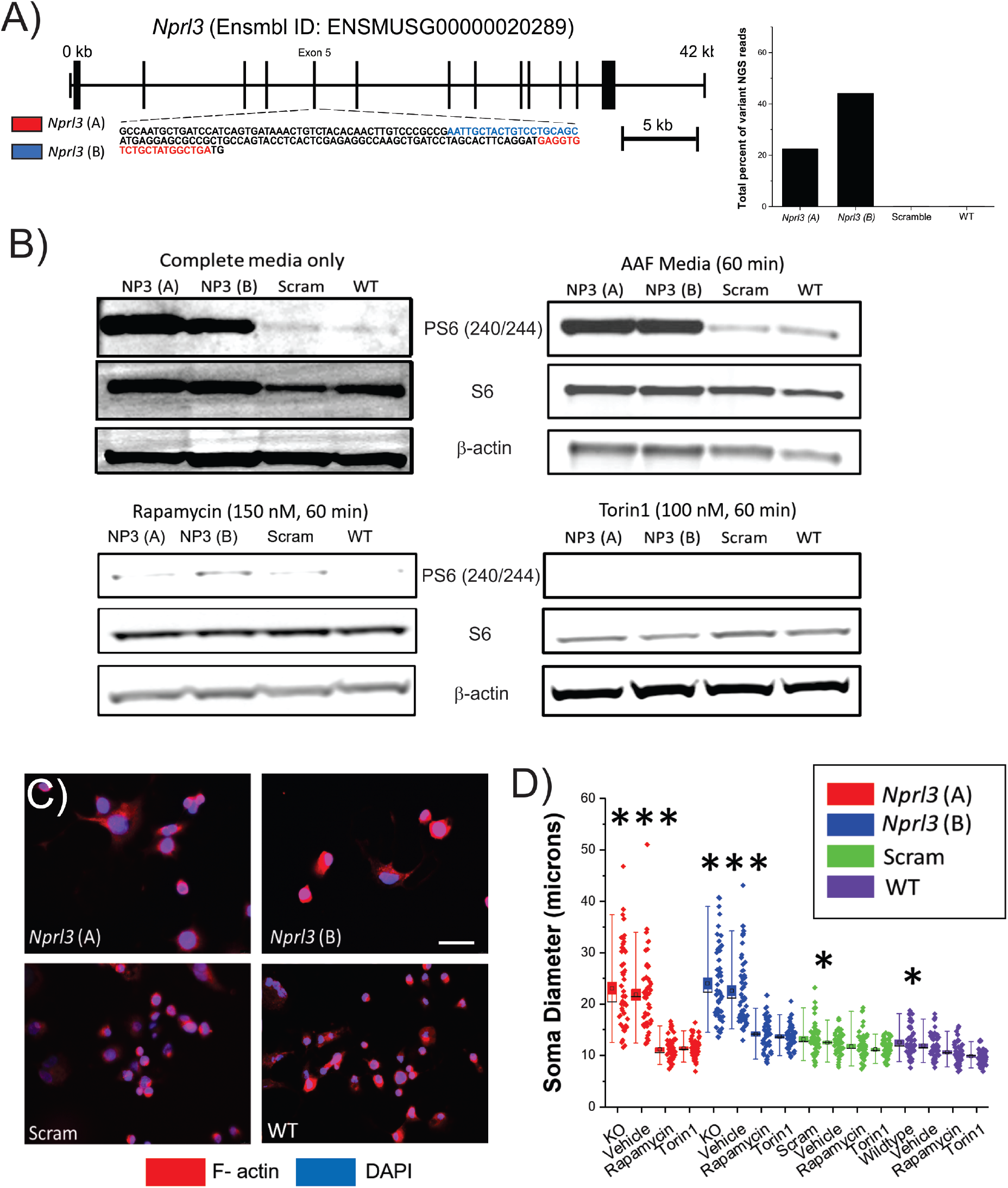
*Nprl3* KO results in mTOR-dependent increases in S6 phosphorylation and soma diameter. The schematic in (A) shows the region targeted by *in silico* generated gRNA. Both gRNA target exon 5 of the mouse *Nprl3* gene. In this schematic, each vertical black bar represents an exon with the thickness of each bar denoting relative size. To define the extend of indel formation, exon 5 PCR products in were sent for whole amplicon next-generation sequencing which revealed abnormal sequences in 22.4% and 40% of NGS reads for amplicons from *Nprl3 (A)* and *Nprl3* (B) lines, respectively (*graph* in A; see Supp. alignment tables). Western assay of *Nprl3* KO N2aC reveals an increase in PS6 (240/244) (B) in complete media that is reversible with rapamycin (150 nM, 60 min) or torin1 (100 nM; 60 min). Sustained levels of PS6 (240/244) were also observed in *Nprl3* KO N2aC after incubation in AAF media vs. scramble and WT cells suggesting that the GATOR1 complex is nonfunctional (AAF media, 60 min). Western blot densitometry data are in Supp. Fig. 1 (n=3 blots per group). Increase in S6 levels is a consequence of activated mTOR signaling and protein translation. In (C), representative images of each cell line are shown. After *Nprl3* KO, a statistically significant increase in soma diameter was observed in both *Nprl3* KO N2aC lines vs. WT and scramble controls but not vehicle (DMSO) treated cells (D, n=50 cells per group, p<0.001). Application of rapamycin (50 nM, 48hours) or torin1 (50 nM, 48 hours) resulted in a statistically significant decrease in soma diameter in all groups, although the maximal reduction was in *Nprl3* KO N2aC. In (D), each box represents the S.E. with a mean line shown, whiskers represent 5-95% confidence intervals, diamonds to the right represent individual data points. ***= p<0.001 vs. scram WT and torin1 or rapa, *=p<0.05 vs torin1 and rapa, WT= wildtype, Scram= scramble, F-actin = filamentous actin, KO= untreated KO, NP3= *Nprl3,* B-actin = beta actin. Calibration mark in (C) = 25 μM

mTOR activation is modulated by cellular amino acid levels via the GATOR1 complex in non-neural cells but this has not been demonstrated in neuronal cell lines. Thus, when incubated in amino acid free (AAF) conditions for 60 minutes, mTOR activation, as measured by S6 phosphorylation levels, was reduced in scramble control and WT N2aC compared with complete media incubation (Fig. 2B; densitometry data in Supp. Fig. 1). In contrast, *Nprl3* KO N2aC incubated in AAF media for 60 minutes exhibited sustained levels of PS6 (240/244), demonstrating that mTOR activation remains enhanced in both complete and AAF media conditions in the setting of *Nprl3* loss and that nutrient deprivation cannot override the effects of *Nprl3* loss on mTOR activation.

### Cell soma diameter

Dysmorphic neurons observed in human FCD II tissue samples resected from patients with known MPG variants are often twice the size of neurons in control brain tissue ^15,23^. Therefore, we hypothesized that *Nprl3* KO in N2aC would result in increased soma diameter. Indeed, a 2-fold increase (p<0.001) in soma diameter was observed in *Nprl3* KO cells visualized with filamentous actin under fluorescence microscopy compared to control cells (25 cells per group; 2 biological replicates Fig. 2C,D). When treated with rapamycin (50 nM) or torin1 (50 nM) for 24 hours, these N2aC showed a reduction in soma diameter (Fig. 2C,D) equivalent to WT cell diameter (p<0.001) demonstrating that *Nprl3* loss resulted in mTOR-dependent soma enlargement^14,15^.

### Constitutive 4E-BP1 phosphorylation during amino acid starvation

In non-neural cell types, mTORC1 is active as a kinase when tethered to the lysosome via RAGs proteins binding partners. To assay lysosomal mTORC1 activity in neurons, we transfected a CFP/YFP FRET biosensor (TORCAR; m**TORC**1 **A**ctivity **R**eporter), that targets the lysosomal membrane by LAMP1 (a lysosomal membrane protein), coupled to 4E-BP1, a direct mTORC1 substrate,^25,26^ to define dynamic changes in 4E-BP1 phosphorylation at the lysosomal surface in living cells. The ratio of CFP to YFP (C:Y) fluorescence (increases or decreases) in transfected *Nprl3* KO lines reflects dynamic changes in C:Y ratio, indicating an increase or decrease, respectively, in 4E-BP1 phosphorylation, and serves as a proxy for mTORC1 activation.

In *Nprl3* KO and control N2aC visualized in complete media for one hour, phospho-4E-BP1 levels were stable as evidenced by non-significant fluctuations in C:Y during this time epoch (Supp. Fig. 3). Next, we evaluated direct mTOR inhibition on 4E-BP1 phosphorylation using torin1 (100 nM) in complete media (CM) as 4E-BP1 phosphorylation has been shown to be relatively unaffected by rapamycin^27,28^. The C:Y ratio was decreased by 5.4% (WT), 6.3% (scramble), 7.5% (*Nprl3* (A)), 5.7% (*Nprl3* (B)) after incubation in torin1 (p<0.001; Supp. Fig. 4). Cells were then incubated in AAF media for 50 minutes after a 10-minute baseline in complete media. While WT and scramble controls cells exhibited an appropriate reduction in C:Y in response to AAF conditions (WT, 9.4% and scramble, 5.7%; p<0.001; Fig. 3C,D,G), *Nprl3* KO N2aC lines showed no change in C:Y (Fig. 3E,F,G) and demonstrated persistent and inappropriate 4E-BP1 phosphorylation under AAF conditions. Thus, *Nprl3* KO resulted in sustained and inappropriate 4E-BP1 phosphorylation (similar to effects on S6 phosphorylation, above) even during amino acid starvation.

**Figure 3:**
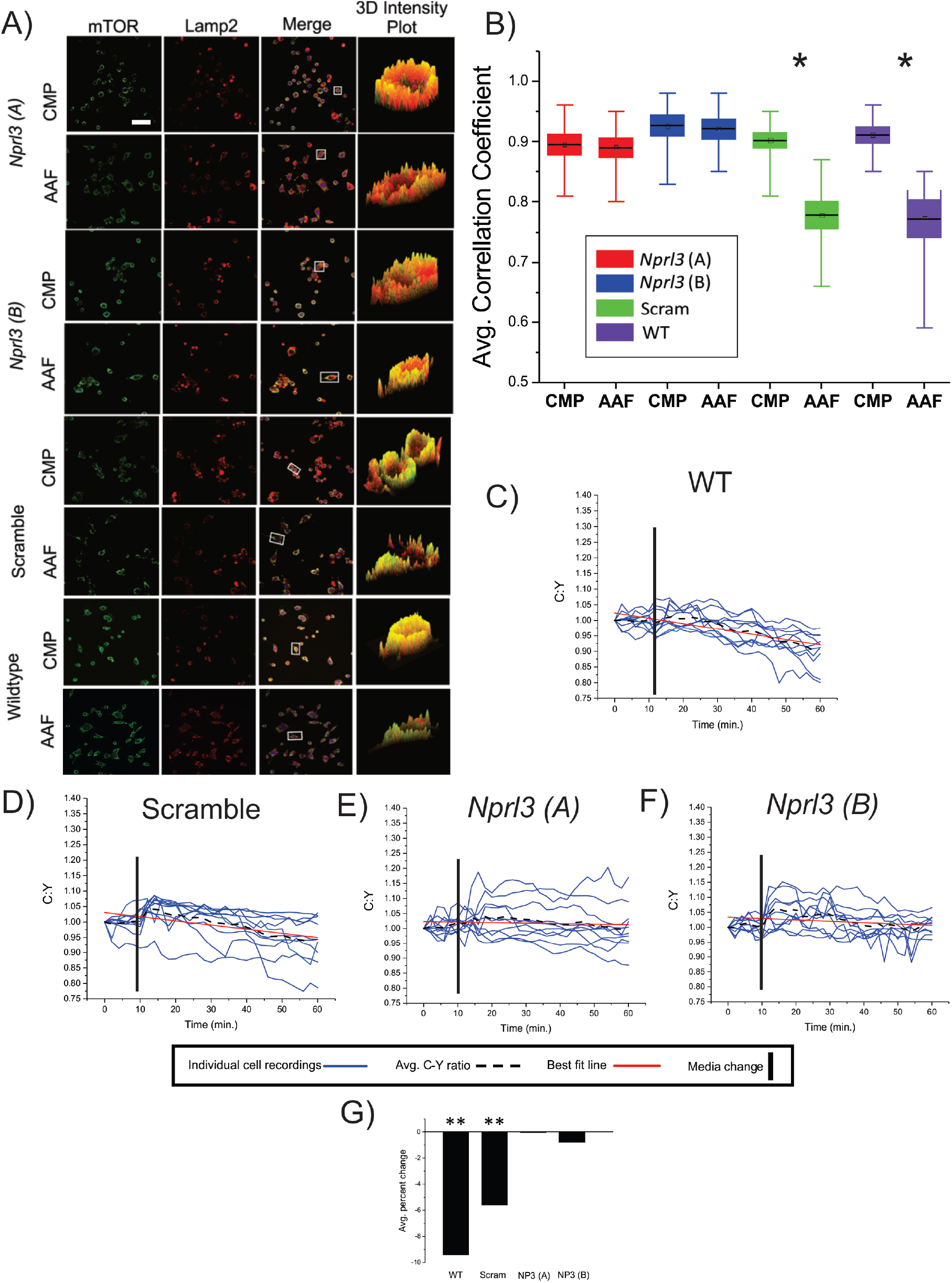
*Nprl3* KO results in persistent mTOR activation at the lysosomal surface in nutrient replete and depleted conditions. ROIs of colocalization between mTOR and LAMP2 are orange or yellow and ROIs where mTOR and LAMP2 are not colocalized appear as distinct red and green areas (A). After incubation in AAF media WT and scramble cell lines display a statistically significant decrease (p <0.05) in colocalization between mTOR and the lysosomal membrane vs. incubation in complete media. However, no statistical difference in colocalization was observed in both *Nprl3* KO cell lines incubated in AAF media vs. complete media. In (B), each box represents the S.E. with a mean line shown, whiskers represent 5-95% confidence intervals. In C-F, C:Y ratio was measured in *Nprl3* KO, scramble control, and WT N2aC lines transfected with TORCAR. Cells were incubated in AAF media for 60 minutes taking measurements at two-minute intervals. A −9.4% and −5.7% decrease in C:Y was observed in WT and scramble N2aC lines during the recording period (C, D, G; p<0.01). with no change observed in *Nprl3* A/B KO lines, respectively (E, F, G). Statistical significance was determined in comparison to corresponding baseline recordings in Supp. Fig. 3. *=p<0.05, ** = p<0.01, WT= wildtype, scram= scramble, NP3 (A) = *Nprl3* A, NP3 (B)= *Nprl3* (B), AAF = amino acid free media, CMP = complete media, Calibration mark in (A) = 25 μM.

**Figure 4:**
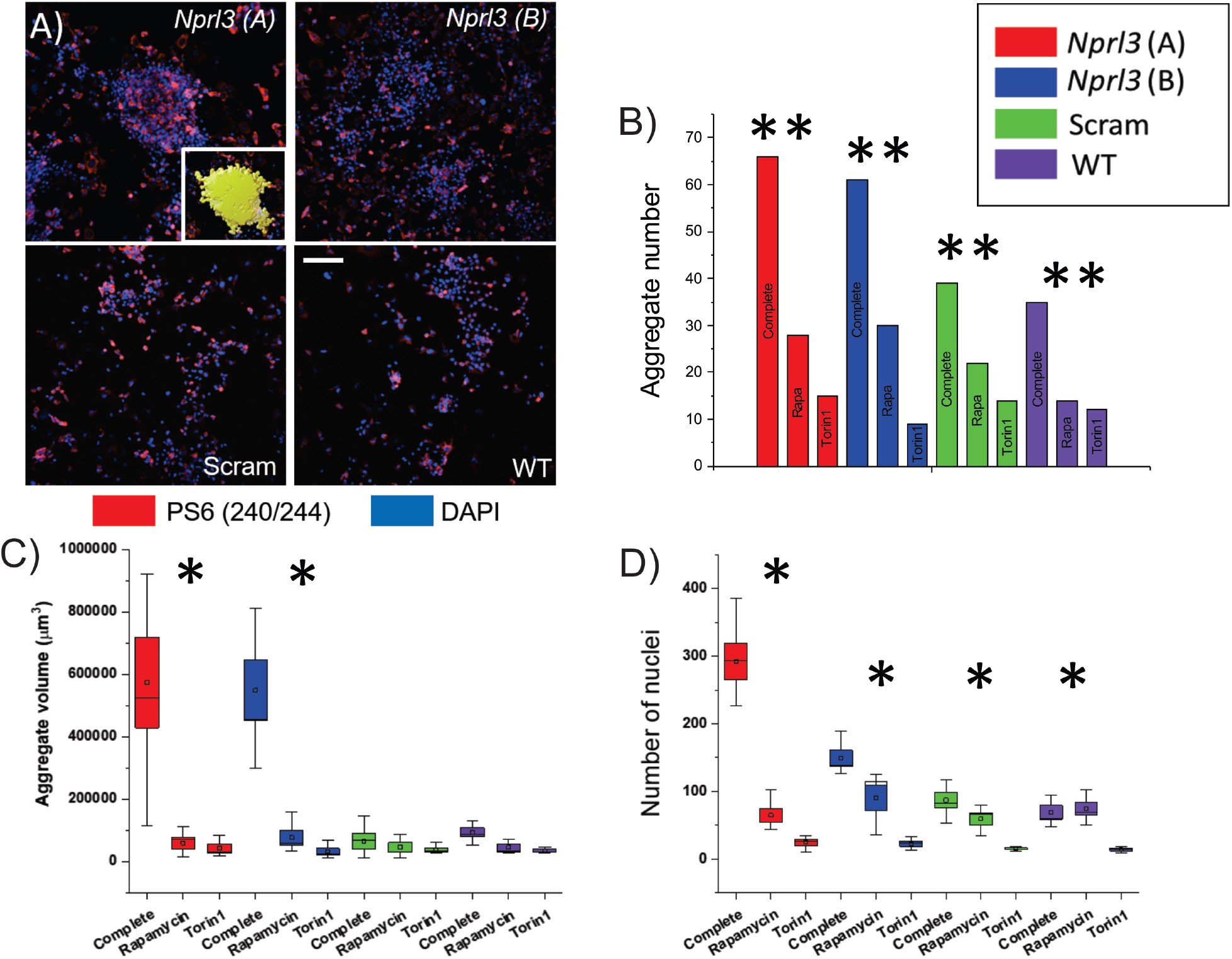
*Nprl3* KO results in mTOR-dependent cellular aggregates *in vitro.* Representative confocal photomicrographs (A) reveal large cellular aggregates in both *Nprl3* KO but not WT or scramble control N2aC lines. Supp. Movie 1 shows time-lapse growth of aggregates in *Nprl3* KO, scramble, and WT cells. *Nprl3* KO N2aC have significantly increased total aggregate counts in digital images taken at low magnification of equal size (B; p< 0.01). The volume of each aggregate (n=5 aggregates per group) is greater in both *Nprl3* KO N2aC lines than in WT or scramble control cells (C; inset in A shows digital surface created to measure aggregate volume; p< 0.05). *Nprl3* KO aggregates have a greater cell number than scramble or WT control cells as measured by DAPI fluorescence (p < 0.05). A proliferation assay did not show an increase in proliferative capacity between *Nprl3* KO and WT or scramble cell lines (Supp. Fig. S2). Large aggregate number, volume, and individual cell number were all preventable with rapamycin (50 nM; 48 hr.) or torin1 (50 nM; 48 hr.) treatment (B,C,D) which was also observed in time-lapse imaging (Supp. movies 2-3) **= p<0.01, *=p<0.05, scram = scramble, WT= wildtype, complete = complete physiological media; rapa= rapamycin. Calibration mark in (A) = 100 μM.

### Nprl3 KO alters lysosomal localization of mTOR

In non-neural cell types, GATOR1 triggers translocation of mTORC1 from the lysosome to the cytoplasm in an inactive conformation^12^ under amino acid depleted conditions. Based on our data that AAF conditions do not diminish S6 or 4E-BP1 phosphorylation, we assessed whether *Nprl3* KO would result in persistent and inappropriate localization of mTOR on the lysosomal membrane under AAF conditions. N2aC were incubated in AAF or complete media for 60 minutes, fixed, fluorescently labeled with anti-mTOR and anti-LAMP2 antibodies, and imaged on a spinning-disk confocal microscope to define the correlation between mTOR fluorescence intensity and LAMP2 fluorescence intensity (Pearson’s coefficient). As shown in Fig. 3A, the orange color corresponds to colocalization between mTOR (green fluorescence) and LAMP2 (red fluorescence) and represents the highest protein colocalization. In contrast, punctate areas of green (mTOR) or red (LAMP2) represent low degrees of colocalization. In complete media, mTOR:LAMP2 colocalization was similar between *Nprl3* KO, scramble control, and wildtype N2aC (Fig. 3B) with R values at or above 0.9 for each group (n=10 per group). However, after incubation in AAF media, mTOR and LAMP2 remained colocalized in Nprl3 KO cells (colocalization R values of 0.89 and 0.92 for *Nprl3 (A)* and *Nprl3 (B)*, respectively) whereas AAF media led to dissociation of mTOR from the lysosomal surface in WT and scramble control cells (p<0.05; n=10 per group; R values of 0.78 and 0.77, respectively, Fig. 3B). In the setting of *Nprl3* loss, mTOR remains co-localized to the lysosomal membrane and activated (evidenced by persistent phosphorylation of S6 and 4E-BP1) even under AAF conditions, demonstrating that *Nprl3* loss decouples mTOR activation from modulation by metabolic state via GATOR1.

### Nprl3 KO results in mTOR-dependent cellular aggregation

Laminar disorganization of the cerebral cortex in human FCD II specimens often includes cytomegalic dysmorphic neurons observed in close physical apposition (clumps or clusters of neurons)^29^ and cellular aggregates have been reported *in vitro* in astrocytes lacking *Tsc1*, a known mTOR modulator^30^. We thus hypothesized that *Nprl3* KO would result in abnormal cellular aggregation *in vitro.* Aggregate formation in live cells was visualized by time-lapse imaging. *Nprl3 KO* N2aC lines formed large aggregates (clumps) that moved as a single unit and accumulated cells from the surrounding environment during the 48-hour imaging epoch. Cells were observed readily adhering to aggregates, but cells were not observed detaching from aggregates (Supp. Movie 1). Aggregation in time-lapse videos was prevented with rapamycin (50 nM) or torin1 (50 nM; Supp. Movies 2 and 3) treatment for 48-hours and during treatment, cells were observed moving freely within the recorded region. In control lines, aggregates were observed rarely and were small in size (Supp. Movie 1; Fig. 4)

We next assayed the morphology of cellular aggregates. Fixed N2aC were probed with PS6 (Ser240/244) antibody and imaged on a spinning-disk confocal microscope. Digital micrographs were reconstructed in 3D to define total aggregate number, cell number within aggregates, and aggregate volume in *Nprl3* KO, scramble, and WT N2aC lines. There was a statistically significant increase in the total number of cell aggregates in *Nprl3* KO N2aC lines (>60 aggregates per region) compared to control lines (<40 aggregates per region; Fig. 4B). Aggregate volume (n=10 per group) was defined in Bitplane using the surface tool (*inset* in Fig. 4A). *Nprl3* KO N2aC lines had statistically significant increases in cell aggregate volume compared to scramble and WT lines (Fig. 4C). Quantifying the number of nuclei in each aggregate (n=10 per group) revealed statistically significant increases in cell number in *Nprl3* KO N2aC lines versus scrambled and WT controls (Fig. 4D). Increases in aggregate number, volume, and increases in cell number per aggregate in *Nprl3* KO cells were prevented with either rapamycin (50 nM) or torin1 (50 nM) for 48 hours during imaging suggesting that cell aggregation due to *Nprl3* loss were mTOR-dependent.

Aggregate formation was not due to enhanced cell proliferation as assayed by EdU (Supp. Fig. 2) or alterations in cell death via assays for necrosis (propidium iodide) and apoptosis (annexin V; Supp. Fig. 2).

### Nprl3 KO in vivo results in mTOR-dependent defects in cerebral cortical lamination

Resected human brain tissue specimens from individuals with *NPRL3* variants show FCD II^31^, including disorganized laminar structure, enlarged dysmorphic neurons ^23^, and heterotopic neurons in white matter (Fig. 5D). To model the effect of *NPRL3* gene inactivation in the developing human brain, we used *in utero* electroporation (IUE) to transfect *Nprl3* (A) and (B) gRNAs and Cas9/GFP plasmids into neuroglial progenitor cells in the telencephalic ventricular zone on embryonic day (E)14. Dual gRNA transfections were performed to assure full *Nprl3* KO. Pups were euthanized on P3 and brain sections were probed with anti-GFP antibodies to visualize cells with *Nprl3* KO. Anti-CTIP2 antibody labeling was used to define layer IV-VI neurons.

**Figure 5:**
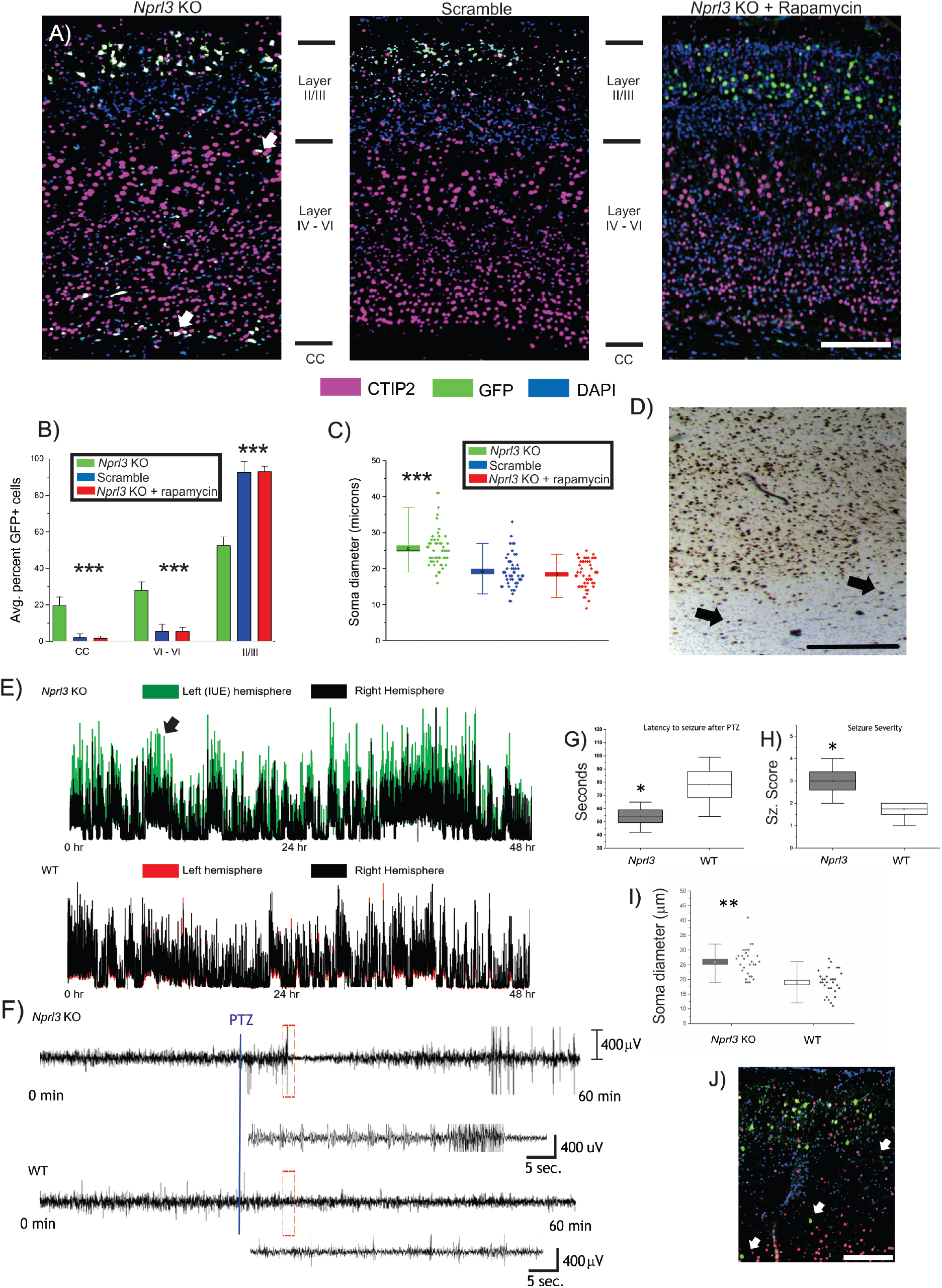
Focal KO *of Nprl3 in vivo r*esults in laminar defects, increased soma diameter, increased cortical excitability, and reduce seizure threshold. At P3, GFP+ cells were observed in the white matter of the corpus callosum and in layers IV-VI in *Nprl3* KO brains but not in scramble control specimens (n=5 per groups; p<0.001) whereas all cells were observed in the birthdate appropriate layers II/III (A,B). GFP+ neurons in *Nprl3* KO specimens did not express CTIP2 indicating that GFP-labeled cells in these layers had failed to attain their appropriate laminar destination in layers II/III. In addition, GFP+ cells in layer II/III in *Nprl3* KO pups were larger than scramble cells in the same layer. Rapamycin treatment prevented laminar defects (A,B) and soma diameter increases (C). In (D), a cortical section stained with NeuN from a surgical epilepsy FCD specimen from a patient in Fig. 1 (subject 18) shows ectopic white matter neurons, similar to what was observed in (A). In (C), each box represents the S.E. with a mean line shown, whiskers represent 5-95% confidence intervals. Five weeks after IUE, mice were implanted with dural EEG electrodes and recorded for 48 hours. In (E), a representative line length analysis of the 48 hr. EEG recording shows increase line length (a proxy for spike amplitude) in the electroporated cortex suggesting a hyperexcitable cortex (n=6 per group) compared to the contralateral cortex and WT mouse cortices where line lengths in the right and left hemispheres are similar. WT or *Nprl3* IUE mice were treated with 55 mg/kg PTZ and EEGs were recorded. Representative EEG tracings are shown in (F). After PTZ injection, *Nprl3* IUE mice had a statistically significant average decrease (p<0.05) in latency to seizure (G; 54.25 sec.) compared to WT mice (G; 78.25 sec.). *Nprl3* KO mice also had increased average seizure severity (SS = 3) compared to WT mice (SS = 1.75) as measured by behavioral seizure scale (n= 6 per group; p<0.05; H). Histology of adult IUE mice implanted with EEGs (J) revealed larger soma diameter in layer II compared to WT MAP2+ neurons (data not shown; p<0.01). **= p<0.01. Calibration mark in (A) = 200 μM; (J) = 500 μM. Sz. = seizure.

All GFP-labeled neurons were observed in layer II in scramble gRNA electroporated P3 pups consistent with cell fate destination to layer II-III for progenitor cells born at E14. In contrast, numerous GFP-labeled neurons were observed in cortical layers IV-VI and in the subcortical white matter in *Nprl3* KO P3 mouse pups (n=5) in addition to scattered GFP-labeled neurons in layer II (Fig. 5A). No GFP-labeled neurons in *Nprl3* KO pups co-expressed CTIP2 indicating that GFP-labeled neurons observed in layer IV-VI were not in fact derived from cortical layer IV-VI precursor cells and that GFP-labeled cells in these layers had failed to attain their appropriate laminar destination in layers II-III. GFP-labeled neurons in *Nprl3* KO pups were larger in layers II-III than corresponding neurons in the same layer in scramble transfected pups (p<0.0001; Fig. 5A,C) commensurate with observations following *Nprl3* KO *in vitro*.

A key pre-clinical question was whether cell size increase and cortical dyslamination caused by *Nprl3* KO were mTOR-dependent. To test this, pregnant dams were treated with rapamycin (1.0 mg/kg; one dose IP) 24 hours after IUE surgery (approximately E15). At P3, histopathological examination revealed normalization of cell size in all GFP-labeled cells to WT neuronal sizes. The laminar defects in P3 pup brain sections were rescued with rapamycin with the preponderance of GFP-labeled cells observed in the appropriate layer II-III position following rapamycin administration on E15 (Fig. 5B, C).

### *Increase cortical excitability and decrease seizure threshold after Nprl3* KO

Mice electroporated with both *Nprl3* (A) and (B) plasmids were implanted with dural electrodes placed adjacent to electroporated cortical regions at 5 weeks of age. During 48 hours of continuous recording, no spontaneous seizures were observed in *Nprl3* KO or WT mice (n= 6 per group). Line length analysis, a proxy for spike amplitude, revealed increased spike amplitude in the electroporated cortex of *Nprl3* KO mice compared to scramble control and WT mice (representative tracing in Fig. 5E). In the line length analysis, high amplitude lines (Fig. 5E; *arrow*) in the electroporated cortex did not correlate with high amplitude line in the contralateral cortex. Thus, high amplitude cortical activity in the electroporated cortex occurred independently of activity in the contralateral cortex. This was not observed in WT mice where left and right hemispheres were synchronous throughout the 48 hour recording (Fig. 5E).

Next, to assay seizure threshold, *Nprl3* KO and WT mice were injected with the pro-convulsant pentylenetetrazol (PTZ; 55 mg/kg, a minimally convulsive dose in WT CD-1 mice ^32^). *Nprl3* KO mice displayed an average decrease in seizure latency seizure from time of PTZ injection (54.25 sec; SEM= 4.8 sec.) compared to WT mice (78.25 sec.; SEM= 9.8 sec; p<0.05;Fig. 5G). In addition, blinded behavioral seizure scoring revealed an average increase in seizure severity in *Nprl3* KO mice (SS = 3) compared to WT mice (SS= 1.75; p<0.05; Fig. 5H). Behavioral seizure scores corresponded with the presence of high amplitude spikes and seizures recorded on EEG. Immunohistochemistry from EEG implanted KO mice at 5 weeks post-IUE (n= 6) revealed a persistent lamination defect with GFP-labeled neurons in cortical layers IV-VI similar to what was observed in P3 pups. In addition, layer II neurons in adult KO mice were larger than MAP2 positive neurons in layer II in WT EEG implanted mice (Fig. 5I,J).

## Discussion

We show that *Nprl3* KO *in vitro* causes in constitutive mTOR signaling activation in nutritionally replete and amino acid free conditions, an mTOR-dependent increase in soma diameter, abnormal cellular aggregation *in vitro,* and inappropriate and persistent colocalization of mTOR to the lysosomal membrane under amino acid starvation. Interestingly, our results with *Nprl3* are similar to what has been observed in patients with variants in *DEPDC5* and mouse models of *Depdc5* loss ^16,17^ suggesting a mechanistic association between *DEPDC5* and *NPRL3* phenotypes, known protein binding partners within GATOR1.

We present the largest and oldest assembled *NPRL3* pedigree reported to date. This cohort of 133 people who share a founder mutation in *NPRL3* (c.349delG, p.Glu117LysFS) that originated with an Old Order Mennonite couple in the 1720’s. As has been reported in non-Mennonite individuals expressing *NPRL3* variants ^21^, there is a wide spectrum of clinical seizure phenotypes, EEG features, and MRI findings, despite a common genotype in our founder population. For example, some *NPRL3* c.349delG heterozygotes develop seizures, while others will not (overall, seizure penetrance was estimated at 28%). In patients with epilepsy, MRI findings revealed a spectrum ranging from no visible abnormalities to hemimegalencephaly, ours being the second reported case of hemimegalencephaly associated with a pathogenic *NPRL3* variant ^9^. Surprisingly, the most severe seizure phenotypes were not always observed in patients with the most abnormal MRIs and in some individuals with seizures, brain MRI was normal. Our belief is that these individuals may have focal MCD too small to detect by clinical MRI examination since the majority of published *NPRL3*-associated cases have been associated with MCD.

As a strategy to understand phenotypic variability, we reasoned that the large pedigree with common Mennonite haplotype background provided a unique opportunity to assess the effects of a gene modifier as, to date, no study has assessed gene modifiers as a cause for clinical heterogeneity in MCD and epilepsy. In our cohort, no single common variant explained the differences in epilepsy penetrance and severity in the *NPRL3* c.349delG cohort. We acknowledge that multiple variants may act independently in each individual or sibship to modify clinical phenotype. An alternate possibility is that phenotypic variability in our cohort and in other epilepsies associated with MPG variants may reflect tissue variant mosaicism i.e., low versus high allelic burden, rather than variant subtype or gene modifier, as we and others have previously suggested^3,4^. Finally, in contrast to *DEPDC5* variants, the Mennonite *NPRL3* variant does not appear to be strongly associated with SUDEP as there were no cases of possible or confirmed SUDEP within this large cohort ^7^. Indeed, while sudden death was a phenotype observed in *Depdc5-*SynCre mice^17^, early death was not a feature in our mice after focal *Nprl3* KO.

GATOR1 component gene variants may be permissive to mTOR pathway hyperactivation by allowing inappropriate and constitutive localization of mTORC1 to the lysosomal membrane, in an active conformation, irrespective of cellular metabolic conditions and even when metabolic cellular demands require mTOR inhibition ^10,15^. In our study, CRISPR/Cas9 *KO* of *Nprl3* resulted in persistent and inappropriate localization of mTOR to the lysosomal membrane during amino acid starvation. Indeed, loss of Depdc5 in mouse renders GATOR1 non-functional^14^ and previous studies have shown that shRNA mediated knockdown of *Depdc5* or *Nprl3* resulted in increased localization of mTOR to the lysosomal surface even during amino acid starvation ^15^.

*Nprl3* KO led to cellular aggregates *in vitro* that were prevented with rapamycin. These aggregates were not the result of increased cell proliferation nor were they due to diminished cell death. Close cellular apposition is a common finding in human brain tissue specimens from patients with FCD II^29^ and mirror previous *in vitro* findings in astrocytes generated from the *Tsc1*-GFAP-cre mouse strain^30^. Cell enlargement and heterotopic neurons in white matter are seen in human tissue specimens from patients with *NPRL3* variants and following *in utero* focal *Nprl3* KO. Interestingly, our results are similar to what has been observed in brain specimens resected from individuals with *DEPDC5* variants^32^ and following *Depdc5* KO *in vivo*^16,17^, suggesting a mechanistic association between *DEPDC5* and *NPRL3* phenotypes; loss of either DEPDC5 or NPRL3, known protein binding partners and functional components within GATOR1, leads to similar, though not identical, phenotypes. *Nprl3* KO did not result in spontaneous electrographic or behavioral seizures as has been observed in a conditional mouse *Depdc5* KO strain or following *Depdc5* KO in rat^16,17^, although a conditional KO of *Nprl3* (versus IUE) and strain differences (rat versus mouse) could account for these disparities. However, we did observe an increase in spike amplitude in the electroporated hemisphere compared to the contralateral hemisphere during baseline recordings and a decrease in seizure threshold of each *Nprl3* KO mouse.

Together, our data provide strong links between mTOR pathway hyperactivation, morphological features, and clinical phenotype caused by NPRL3 variants. We show that mTOR inhibition can rescue these changes and thus, these findings provide a platform for studies examining the efficacy of mTOR inhibitors for the treatment of seizures in patients with GATOR1 gene variants.

## Methods

### Collection and characterization of NPRL3 variants in the Amish and Mennonite Communities

*NPRL3* mutation heterozygotes (n=133; c.349delG, p.Glu117LysFS) were identified from the Mennonite community between 2016 and 2020 through whole exome sequencing, targeted *NPRL3* variant testing, the Invitae Epilepsy Panel (Invitae Corporation, San Francisco, CA) or the Plain Insight Panel, an extended carrier test developed for the Old Order Amish and Mennonite communities. Cascade testing of extended family members was used to identify further potential heterozygotes. Seizure phenotypes were determined through structured medical interviews with parents of affected children (<18 years old) and adult subjects (≥18 years old). While 48 subjects *NPRL3* mutation heterozygotes reported having seizures, 6 were excluded from the structured medical interview due to a dual diagnosis. All patients or their parents were consented for the survey over the phone or in writing. When available, MRI and EEG reports were collected and analyzed. MRIs and EEGs from three subjects were excluded from analysis due to a dual diagnoses.

Genome-wide association analyses were performed with whole exome sequencing data generated at the Regeneron Genetics Center. Individual vcf files for each *NPRL3* c.349delG heterozygote (n=83) were parsed using FileMaker Pro (FMP) scripts and uploaded into a custom FMP database housing >4000 Amish and Mennonite exomes. Variants were annotated with SnpEff and ANNOVAR. The variant data (n=230,656) were then filtered based on genotype quality (>98), allele balance (<3), minor allele frequency (>0.1), and a consistency check for Hardy-Weinberg equilibrium. Surviving variants (n=71,199) were then analyzed with a chi-square statistic to assess differences in minor allele frequencies in the *NPRL3* c.349delG cohort versus 835 Old Order Mennonite controls.

### Exome sequencing

Exome sequencing was performed in collaboration with the Regeneron Genetics Center (RGC) as previously described^38^. In brief, 1μg of high-quality genomic DNA was used to prepare exome-captured libraries using the NimbleGen VCRome SeqCap 2.1 or IDT XGen targeted capture reagent. Libraries were sequenced on the Illumina HiSeq 2500 or the NovaSeq 6000 sequencing platforms achieving coverage of >85% of bases at 20x or greater. Raw sequence reads were mapped and aligned to the GRCh38/hg38 human genome reference assembly using a cloud-based bioinformatics production pipeline that uses BWA-mem for mapping and alignment and GATK HaplotypeCaller for variant and genotype calling. Genomic variants were annotated using an RGC implemented analysis and annotation pipeline that uses Annovar and customized Perl scripts.

### Collection, processing, and analysis of human tissue specimens from patients with NPRL3 variants

Three patients underwent brain resection to treat intractable epilepsy and FFPE blocks and slides of preserved brain tissue were obtained with patient consent (McGill University Health Center-Glen Site Department of Pathology, Children’s Hospital of Akron Department of Pathology & Laboratory Medicine, The Children’s Hospital of Philadelphia Department of Pathology and Laboratory Medicine).

### CRISPR/Cas9 construct generation and validation

Guide RNA (gRNA) targeting the spCas9 endonuclease to regions in the mouse genome encoding *Nprl3* were calculated *in silico* using ChopChop software (chopchop.cbu.uib.no). Two gRNA (*Nprl3* (A) and *Nprl3* (B)) with sequences -GAGGTGTCTGCTATGGCTGA- and - AATTGCTACTGTCCTGCAGC-targeting exon 5 of *Nprl3* were used for these experiments (Fig. 2A). A scramble gRNA (-GACTACCAGAGCTAACTCA-) was used as a transfection and gRNA control. gRNAs were then assembled into oligonucleotides (Integrated DNA Technologies, Coralville, IA), annealed using ligase buffer (Promega, Madison, WI) at 98ºC for 5 min.

To validate that our gRNA containing CRISPR/Cas9 plasmid created indels in our regions of interest, DNA from *Nprl3* or and scramble FAC-sorted cells lines (as described below) as well as wildtype (WT) N2aC were assayed for mis-matched DNA pairs via T7 endonuclease assay (EnGen Mutation Detection Kit; New England Biolabs, Ipswich, MA) with PCR primers targeted towards our genomic region of interest (Integrated DNA technologies, Coralville, IA). To define the sequence misalignments generated by Cas9 editing of *Nprl3,* PCR amplicons of exon 5 of the mouse *Nprl3* gene were purified using a PCR purifications Kit (Qiagen, Hilden, Germany) and sent to the Massachusetts General Hospital Center for Computational and Integrative Biology DNA Core for whole amplicon next-generation sequencing (NGS; Massachusetts General Hospital Center for Computational and Integrative Biology DNA Core). These data revealed double strand breaks and numerous misaligned sequences in *Nprl3 (*A) and *(B)* KO cell lines and no misaligned sequences in WT or scramble lines (Supp. alignment tables).

### Cell Culture and establishment of KO cell lines

Neuro2a cells (N2aC; Sigma-Aldrich, St. Louis, MO) express many neuronal markers, have a high transfection efficiency, and are readily FAC sorted. N2aC were cultured in complete medium consisting of EMEM (Invitrogen, Carlsbad, CA) supplemented with 10% FBS (Invitrogen, Carlsbad, CA). To create stable CRISPR/Cas9 edited cell lines, N2aC were transfected using Lipofectamine LTX with Plus reagent (ThermoFisher Scientific, Waltham, MA) and 30 μg of plasmid diluted in 300 μl Opti-MEM (Invitrogen, Carlsbad, CA) for 48 hours. After 48hours of transfection, cells were trypsinized (0.25%), centrifuged, washed with ice-cold PBS, passed through a cell strainer into a 5 ml conical tube and assayed by flow cytometry (University of Maryland School of Medicine Flow Cytometry Core) for sorting based on mCherry (Cas9) fluorescence (BD FACSAria II cell sorter; Becton Dickinson and Company, Franklin Lakes, NJ). mCherry+ sorted cells were placed into PBS containing 1% serum until re-plating. Cells were re-plated in complete media and grown to confluence.

### Analysis of soma diameter

Cell soma diameter was measured in digital images of CRISPR-edited, scramble control, or WT N2aC lines in ImageJ ^34^. Each measurement was taken using the longest dimension of each soma in 50 total cells (25 cells in 2 biological replicates). To define the mTOR-dependency of cell size changes, cells were incubated with the mTOR inhibitor rapamycin (50 nM, 24hours, Cell Signaling Technologies, Danvers, MA) or torin1, a specific ATP-competitive dual kinase inhibitor of mTORC1 and mTORC2 (50 nM; 24 hours; Torcris, Bristol, UK) using DMSO as a vehicle control. As high concentrations of mTOR inhibitors can results in cell death during long incubation periods, we have used a lower concentration of both rapamycin and torin1 in these experiments to facilitate changes in cell size without inducing cell death. Statistical analysis of cell size differences was assessed in Origin (Northampton, MA) using one-way ANOVA (p < 0.05 considered statistically significant). Measurements were graphed using a box and whisker plot where the box represents the standard error of the mean and the whiskers represent the 5-95% confidence interval.

### Analysis of lysosomal mTORC1 activity using live-cell imaging

*Nprl3* KO, scramble, or WT N2aC were plated on video dishes. Cells were grown to 30-40% confluence and transfected with TORCAR (pcDNA3-Lyso-TORCAR was a gift from J. Zhang; Addgene plasmid # 64929; http://n2t.net/addgene:64929; RRID:Addgene_64929) as described above. After 48 hours, transfection media was removed, and warm complete imaging media was washed on. Cells were cultured for an additional 24 hours in complete DMEM imaging media (FluoroBrite DMEM, ThermoFisher Scientific, Waltham, MA).

TORCAR transfected cells were imaged on a Zeiss LSM Duo slit-scanning confocal microscope (Oberkochen, Germany) with lasers allowing us to visualize CFP and YFP channels simultaneously. Images were taken at 40x magnification recording fluorescence from the CFP and YFP channels using time-lapse imaging at two-minute intervals for 60 minutes. First, a 10-minute baseline in complete imaging media was established and then experimental media was washed on. Experimental conditions were complete media baseline for 60 minutes, AAF imaging media (no phenol red; US Biologicals, Salem, MA), or complete imaging media with torin1 (100 nM). 10 cells were recorded for each experimental condition.

After recording CFP and YFP fluorescence data, the CFP to YFP ratio (C:Y) was calculated, averaged across 10 cells per group and a best-fit line was generated in Origin statistical software (Northampton, MA). For each group, the individual C:Y for each cell was graphed (blue) along with the average C:Y (dashed black line) and best-fit line (red). All data were normalized to account for focal plane changes and differences in transfection efficiency. Percent change was also calculated using the averaged C:Y. Statistical differences were determined using a one-way ANOVA with a p value of < 0.05 deemed significant. Statistical significance was determined using the average C:Y ratio during each 60-minute baseline in complete media and compared to the overall change in C:Y for each experimental manipulation.

### Live-cell Imaging of cellular aggregates

*Nprl3* KO, scramble, and WT N2aC were trypsinized, resuspended, and counted using a digital hemocytometer. Prior to plating cells for experimentation, any pre-existing aggregates were removed by passage through a cell strainer to create a single-cell suspension. To ensure that cellular aggregates were not a result of over-plating, all lines were then plated at equal density into video dishes or chamber slides. Cells were allowed to adhere to video-dishes for 24-hours. After 24-hours, dishes were placed into an Olympus VivaView (Center Valley, PA, USA) imaging incubator maintained at 37° C with 5% CO2. Using the accompanying software, three regions of interest were selected in each of four video dishes-*Nprl3* A, *Nprl3* B, scramble, WT. Images were taken at 10x every 30 minutes for 48-hours. For rapamycin and torin1 treated cells, the experimental procedure above was repeated with the addition of either 50 nM rapamycin or 50 nM torin1 to cell culture media immediately prior to imaging. Time-lapse images were compiled into videos using ImageJ ^34^.

### Immunocytochemistry

N2aC were fixed in 4% PFA at room temperature for 20 minutes and then permeabilized in phosphate-buffered saline (PBS) containing 0.3% Triton X-100 (ThermoFisher Scientific, Waltham, MA). Cells were blocked for 2 hours at room temperature (RT) in 5% normal goat serum (Jackson ImmunoResearch, West Grove, PA). Cells were incubated in one of the following primary antibodies in blocking solution containing 5% normal serum at 4°C overnight: F-actin (1:1000; Abcam, Cambridge, UK), mTOR (1:1000; Abcam, Cambridge, UK), lysosome-associated membrane protein 2 (LAMP2; 1:1000; Thermofisher, Waltham, MA), or phopsho-ribosomal S6 protein (PS6; Ser240/244; rabbit monoclonal; 1:1000; Cell Signaling, Danvers, MA). The secondary antibody, containing a fluorochrome (Alexa Fluor 488 or Alexa Fluor 594; all 1:1000 dilution; Molecular Probes, Eugene, OR), was incubated with the cells for 2 hours at RT. DAPI counterstain was used to visualize nuclei and was applied along with mounting media during cover slipping.

Apoptosis and necrosis assays were performed using live *Nprl3* KO, scramble, or WT N2aC plated in chamber slides using the Annexin V-FITC Early Apoptosis Detection Kit (Cell Signaling Technology, Danvers, MA). As a positive control for apoptosis and necrosis, 25 nM Etoposide was incubated with WT N2aC for 24 hours. As a negative control, WT N2aC were used after incubation in complete media.

A proliferation assay was performed on *Nprl3* KO, scramble, or WT cells using the Click-it Edu assay from Invitrogen (Carlsbad, CA, USA) according to the manufacturer’s instructions. Cells were then fixed in 4% PFA for 20 minutes, cover slipped with DAPI, and imaged on a spinning-disk confocal microscope at 4x.

### Western blot analysis

Cells were lysed in RIPA lysis buffer (50 mM Tris HCl, pH 8.0; 150 mM NaCl; 1% NP-40; 0.5% sodium deoxycholate; and 0.1% SDS) with a protease (Sigma-Aldrich, St. Louis, MO) and phosphatase inhibitor (ThermoFisher Scientific, Waltham, MA) followed by brief sonication. Lysates were centrifuged at 4°C for 20 minutes at 14,000 *g*, and supernatants collected. Protein concentrations (μg/ml) were determined on a nanodrop spectrophotometer (ThermoFisher Scientific, Waltham, MA). A sample mixture containing 30 μg of protein, Nupage Reducing Agent and Nupage Loading Buffer (Invitrogen, Carlsbad, CA) was denatured for 10 minutes at 90°C and loaded onto a Bolt BT Plus 4-12% gel (Invitrogen, Carlsbad, CA). After electrophoresis, proteins were transferred onto PVDF membranes (Immobilon; EMD millipore, Darmstadt, Germany) at 4°C. The membranes were blocked in Odyssey Blocking Buffer (Li-Cor, Lincoln, Nebraska) for 1 hour. at RT. Membranes were probed overnight at 4°C in Odyssey Blocking Buffer with antibodies recognizing PS6 (Ser240/244; rabbit monoclonal; 1:1000; Cell Signaling, Danvers, MA), total S6 ribosomal protein (mouse monoclonal; 1:500; Cell Signaling, Danvers, MA). Odyssey secondary antibodies 800CW or 680RD were used to visualize bands on the blot (1:1000; Li-Cor, Lincoln, Nebraska). To ensure equal protein loading, antibodies recognizing β-actin (1:10,000, Abcam, Cambridge UK) followed by Odyssey secondary (1:1,000, 2-hours, RT; Li-Cor, Lincoln, Nebraska) were used. Blots were developed using an Odyssey Clx scanner and scanned at 169 μm resolution with exposure for both 700 and 800 nm channels set at 3.5 (relative intensity, no units; Li-Cor, Lincoln, Nebraska). All blots were performed in triplicate using distinct biological replicates. Densitometry analysis was performed in Image J ^34^ normalizing band density to β-actin loading control. Densitometry data were analyzed in Origin software (Northampton, MA). Band density from each blot was averaged and the standard error calculated.

### mTOR-lysosomal colocalization analysis

*Nprl3* KO, scramble, and WT N2aC lines were plated into chamber slides and allowed to adhere for 24-hours After 24-hours, cells were incubated in AAF or complete media for 60 minutes and then PFA fixed. Cells were labeled with mTOR and LAMP2 (a lysosomal marker) antibodies as described above. Using a spinning disk confocal microscope (Nikon, Tokyo, Japan), cells were selected and a region of interest (ROI) was isolated. Fluorescence intensity of mTOR and LAMP2 was measured within the ROI using ImageJ ^34^. A Pearson’s correlation test (Excel, Microsoft, Redmond, WA) was then performed to determine whether or not an increase in fluorescence intensity in mTOR correlated with an increase in fluorescence intensity in LAMP2 - the higher the correlation coefficient (R), the greater the degree of colocalization between mTOR and LAMP2. ROIs were measured for 10 N2aC in *Nprl3* KO, scramble control, and WT cells each in each experimental condition. Statistical significance was determined using a one-way ANOVA of averaged R values, with p < 0.05 deemed significant. To better visualize the degree of colocalization in digital images, each ROI was reconstructed into a 3D fluorescence intensity surface plot.

### In utero electroporation

For these experiments, both *Nprl3* (A) and *Nprl3* (B) plasmids were co-electroporated into neural progenitor cells. The two gRNAs were cloned into separate plasmids containing the gRNA and gRNA scaffold via Golden Gate Assembly (courtesy of the Poulopoulos lab, University of Maryland School of Medicine). As each CRISPR “cut” is an independent event in each cell, electroporating two validated gRNA increases the chance of successfully knocking out *Nprl3* in neural progenitor cells. Electroporated cells were identified by GFP expression using one of two strategies: 1) Cas9 linked to GFP via a multicistronic T2A element and preceded by a loxP site and electroporated along with a CRE recombinase plasmid to facilitate GFP expression or 2) Cas9 linked to GFP via a multicistronic element alone (both courtesy of the Poulopoulos lab, University of Maryland School of Medicine). These strategies were equally successful in producing GFP expression in electroporated mouse pups.

Timed-pregnant mice were obtained from Charles River. All procedures were performed on embryonic day 14 (E14). Dams were anesthetized using inhaled isoflurane anesthesia maintained throughout the procedure. Deep anesthesia was confirmed by lack of response to a toe pinch. Once anesthetized, the animal was given an SQ injection of 0.3% buprenorphine for analgesia and placed on a water-circulating heating pad. Hair was removed from the dam’s abdomen using a chemical hair removal agent (Nair, Church and Dwight). A povidone-iodine gel was swabbed over the exposed skin and sterile gauze were placed over the incision site. A 4 cm vertical incision was made along the midline and the uterine horns were exposed using forceps. Uterine horns were washed with warm PBS (37°C) as necessary throughout the procedure to prevent drying of the uterine horns. Using a sharpened microcapillary tube, plasmid DNA was mouth-pipetted into the lateral ventricle of 3-7 embryos per dam, depending on litter size and access to the lateral ventricles. Electrode forceps were place on either side of the embryos head and 5 electrical pulses were passed through the embryo to drive the plasmid into neural progenitors lining the ventricle. Uterine horns were placed back into the dam, the abdominal muscles were sutured closed with 5-gauge silk sutures, and the abdominal skin was closed using veterinary staples. Dams were allowed to recover in a heated cage before returning to normal housing. Dams were given gel food daily and buprenorphine twice daily for 72 hours. post-surgery. One cohort of dams was treated with a single rapamycin injection (1.0 mg/kg) 24 hours post-operation. Mouse pups were born between E20 and 21 and examined for signs of successful electroporation (GFP fluorescence through the skull, burn marks from the electrodes) under a fluorescent dissecting microscope. Pups were euthanized at P3 for IHC analysis or grown for 5 weeks for EEG implantation, seizure threshold testing, and IHC.

### EEG implantation and recording

Implantation and recording were performed using Pinnacle Technology 3 channel EEG system (Lawrence, KS, USA). Implantation was performed using aseptic technique, under isoflurane anesthesia. A rostral-caudal incision was made in the scalp and membranous tissue located under the scalp was cleared away. The skull was dried with ethanol and the implant was secured onto the skull using cyanoacrylate 3 cm behind the bregma. A 23-gauge needle was used to create pilot holes for screws. Each screw was advanced into the skull after silver epoxy was applied to the threads. A novel departure from the standard EEG implantation procedure provided by Pinnacle is the use of flowable light-cured dental composite (Benco Dental, Pittston, PA, USA). Liquid composite was placed over each screw and around the base of the implant to insulate the screws and further secure the device to the skull. Blue light was then used to quickly harden the composite. A light-cured composite, rather than chemically curing resin, was chosen for its biocompatibility, reduction in surgery time and hardening properties that eliminated thermal damage to the mouse from the curing reaction and unnecessary shifts in the EEG implant due to contraction of the composite. Sutures were used to close the incision. Mice were allowed to recover for a minimum of five days. EEG recording is achieved using Pinnacle Technology Seizure Software (Lawrence, KS, USA) and USB-powered EEG connection to the animal. EEGs were recorded for 48 hours. Recorded EEGs were analyzed using Sirenia Seizure Pro (Pinnacle Technology, Lawrence, KS).

### Seizure Threshold Testing

Mice were electroporated as described and maintained in the animal facility for 5 weeks prior to EEG implantation. After a 48-hr. continuous EEG, A 20-minute baseline recording was taken prior to pentylenetetrazol (PTZ) injection. PTZ is a chemoconvulsant commonly used to induce seizures and define seizure threshold in animals where spontaneous seizures are not observed ^35^. Based on an extensive literature search ^32^ we determined that 55 mg/kg PTZ was an appropriate dose to provoke mild seizures (seizure score of 2; see below) in CD1 mice. After a 30-minute baseline, 5-week-old *Nprl3* electroporated mice were given 55 mg/kg PTZ and observed for EEG and behavioral seizures for 40 minutes. A behavioral seizure scale was used to define and quantify the manifestations of seizure in real-time. Seizures were scored as follows: 1) Raised tail and/or abnormal posturing; 2) Myoclonic extension of limbs, favoring one side; 3) Brief tonic-clonic seizures; 4) Tonic-clonic seizures with rearing or jumping; and 5) *status epilepticus* (SE). 30 minutes after PTZ injection, the experiment was terminated, and animals were euthanized using CO2 asphyxiation and thoracotomy. Brain specimens were collected and processed as described below. Data were analyzed in Sirenia Seizure Pro (Pinnacle Technology, Lawrence, KS).

### Immunohistochemistry of animal tissue

Electroporated post-natal day 3 (P3) mouse pups were anesthetized with ice anesthesia. Deep anesthesia was determined by a lack of response to a toe pinch. Adult mice were euthanized by CO2 asphyxiation followed by thoracotomy. All animals were perfused with ice-cold saline and 4% PFA. Brains were dissected out and post-fixed in 4% PFA overnight (12-hours for P3 pups, 24-hours for adult mice). After fixation, brains were embedded in paraffin wax, sliced on a microtome at 8 μm, mounted on slides, and hot air dried overnight. Following drying, brains were deparaffinized, subjected to antigen retrieval via citrate buffer, and washed in distilled water. Washed slides were blocked for 1 hr. at RT in blocking solution containing in 0.3% tween PBS with 5% normal goat serum and incubated in primary antibodies overnight in blocking solution. Primary antibodies used were GFP (1:1000; Abcam), CTIP2 (1:500; Abcam), and MAP2 (1:100; Abcam). After incubation with primary antibodies, slides were washed in PBS and placed in blocking solution with secondary antibodies for 2-hours at RT. Secondary antibodies were Alexa 488, 647, and 594 respectively (1:1000; Invitrogen). After 2-hours, slides were washed in PBS and distilled water and finally coverslipped with Vectashield media containing DAPI.

### Statistics

All statistics were performed in Origin Software. For all analysis, ANOVA was used and the mean and standard error were calculated. For each experiment, statistical significance was determined using a p < 0.05 as the threshold. Statistics for each experiment are detailed in the appropriate section in the methods.

### Study Approval

This study was approved by the University of Maryland School of Medicine Internal Review Board and Institutional Animal Care and Use Committee as well as the Lancaster General Hospital Institutional Review Board

## Supporting information

Nprl3 A seq alignment

Nprl3 B seq alignment

## Author Contributions

PHI – *in vivo* experiments, imaging, data analysis, EEG implantation, seizure threshold testing, EEG analysis, manuscript writing. MEE-human data collection and analysis, manuscript writing. KMCP-human data collection and analysis. LEB-human data collection and analysis. AEB – Cell culture, Western assay, immunocytochemistry, data analysis. JKB-IUE and histology. AR-IUE and histology. MB – Cell culture, Western assay, immunocytochemistry, cell size measurements. EGP-exome and sequencing data analysis, CGJ-exome sequencing. AP-conceptualization, supervision, plasmid design. VJC-human data collection and analysis, supervision. PBC-conceptualization, supervision, data interpretation, manuscript writing.

## Acknowledgements

The authors would like to thank: *NPRL3* patients and their families; Dr. Joseph Mauban, Manager of the University of Maryland School of Medicine Confocal Microscopy core, for his assistance with both live-cell slit-scanning and spinning disk confocal microscopy. Dr. Xiaoxuan Fan, Director of the Flow Cytometry Shared Service at the University of Maryland for his assistance in establishing CRISPR cell lines. Drs. Alex Poulopoulos and Ryan Richardson at the University of Maryland School of medicine for assistance in generating PX330 CRISPR plasmids. Philip H. Iffland, DDS for assistance with improving the EEG implantation technique. This work was supported by NINDS R01NS099452 and R01NS094596-01A1 (PBC). The authors declare no competing interests.

---

**Supp. Figure 1:**
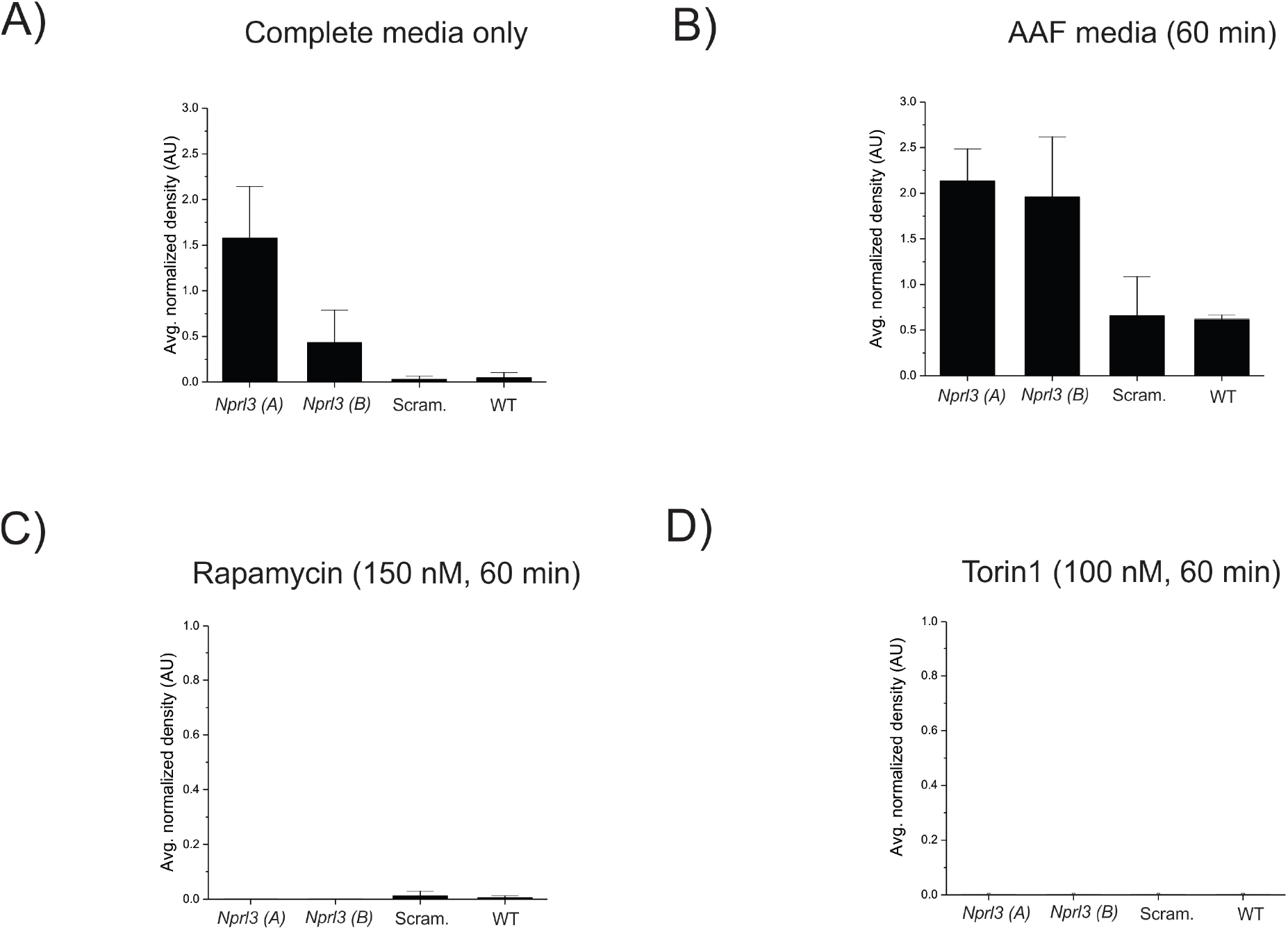
Densitometry data for Western blots in Figure 2. All densitometry measurements are PS6 (240/244) levels normalized to loading control (β-actin) and performed in triplicate using biological replicates. Levels of phosphorylated S6 (240/244) (A-D) were determined using lysates made after incubation in complete or AAF media or complete media containing rapamycin (150 nM) or torin1 (100 nM) for 1 hour. Error bars indicate SEM.

**Supp. Figure 2:**
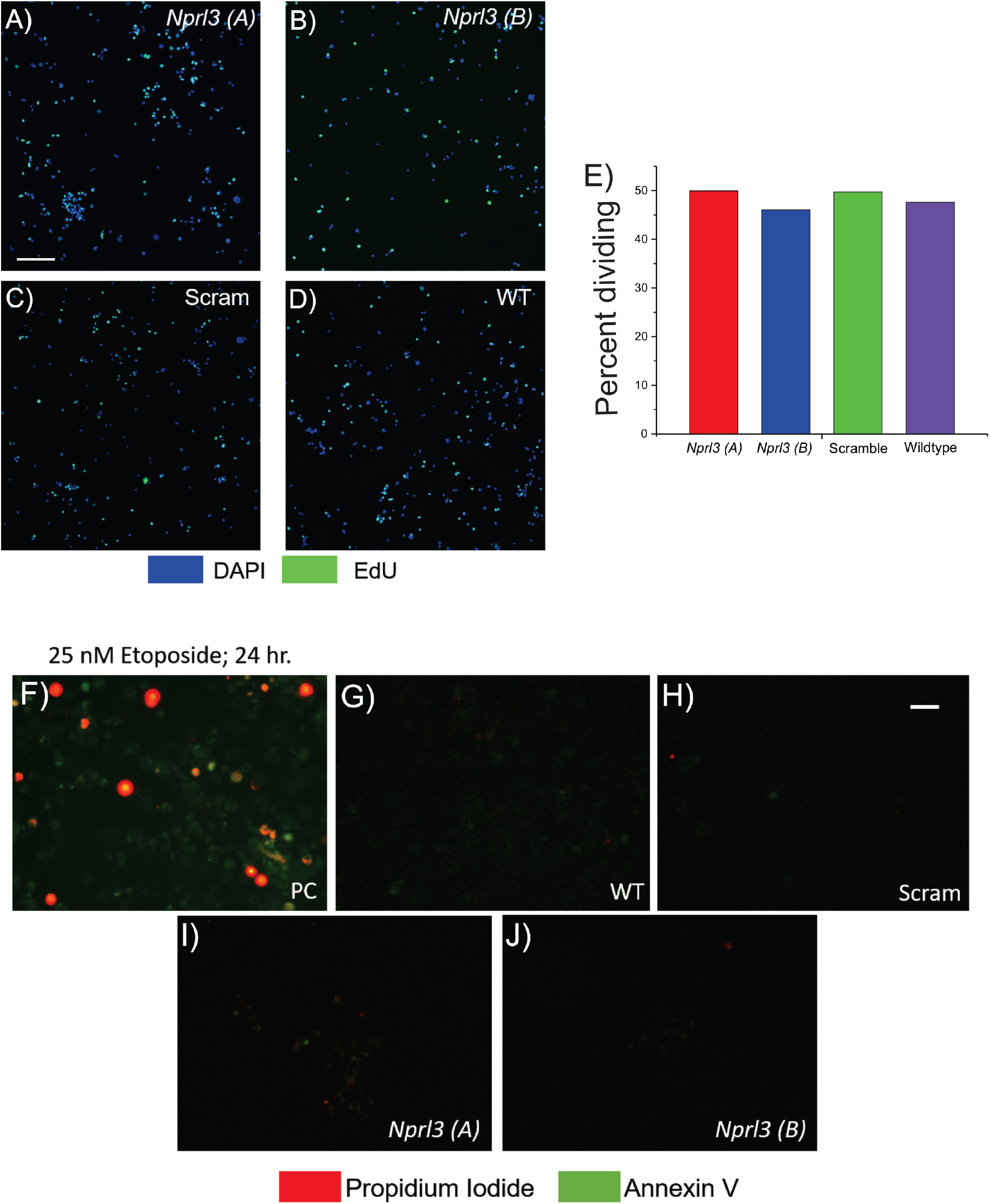
*Nprl3* KO does not enhance cell proliferation or induce cell death. An EdU-based proliferation assay was performed according to the manufacturer’s instructions. Panels A-D show representative images of EdU positive (green; proliferating) and DAPI positive cells. The graph in E shows the percent of proliferating cells in each experimental group (n= 600 cells from two biological replicates of 300 cells each). These data reveal no difference in the percentage of proliferating cells in each group. Micrographs F-J show representative images of cells stained with both propidium iodide, a marker of necrosis (red), and annexin V, an early marker of apoptosis (green). As a positive control (F), WT cells were treated with etoposide for 24 hours. These cells show marked staining for both propidium iodide and annexin V while untreated WT, scramble, and both *Nprl3* KO N2aC do not show high levels of staining for either maker (F-J). Calibration mark in (A) = 500 μM; (H) = 25 μM.

**Supp. Figure 3:**
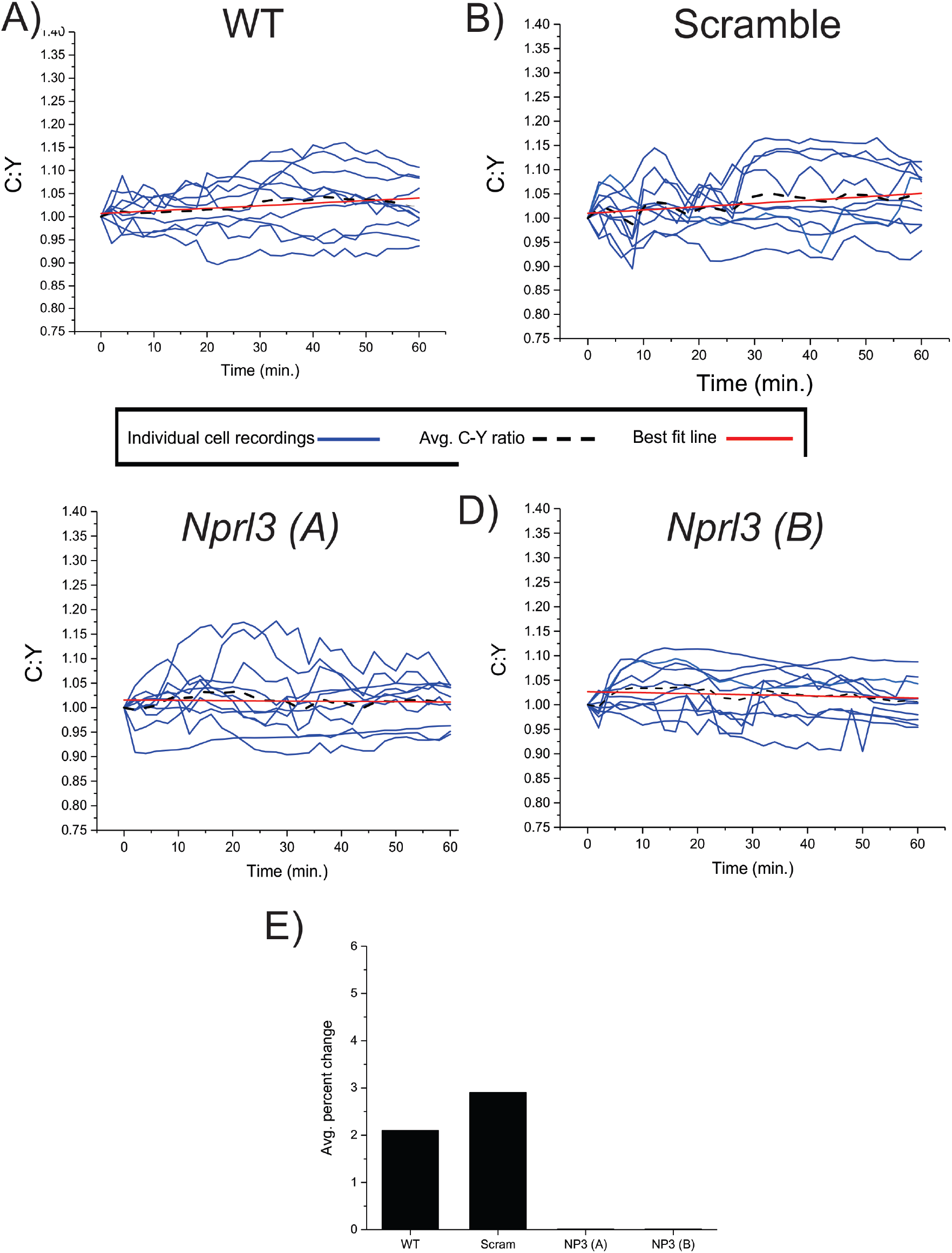
Baseline TORCAR recording show no change in 4E-BP1 phosphorylation. C:Y was measured in *Nprl3* KO, WT and scramble N2aC lines transfected with TORCAR. Cells were incubated in complete media for 60 minutes taking measurements at two-minute intervals. A 2% and 2.8% increase in C:Y was observed in WT and scramble N2aC lines, respectively, during the recording period (A,B). In *Nprl3* KO N2aC a 0.1% increase and a 0.5% increase was observed in *A* and *B* lines, respectively. All statistical calculations in Figs. 3 and S4 were made in comparison to these baseline recordings.

**Supp. Figure 4:**
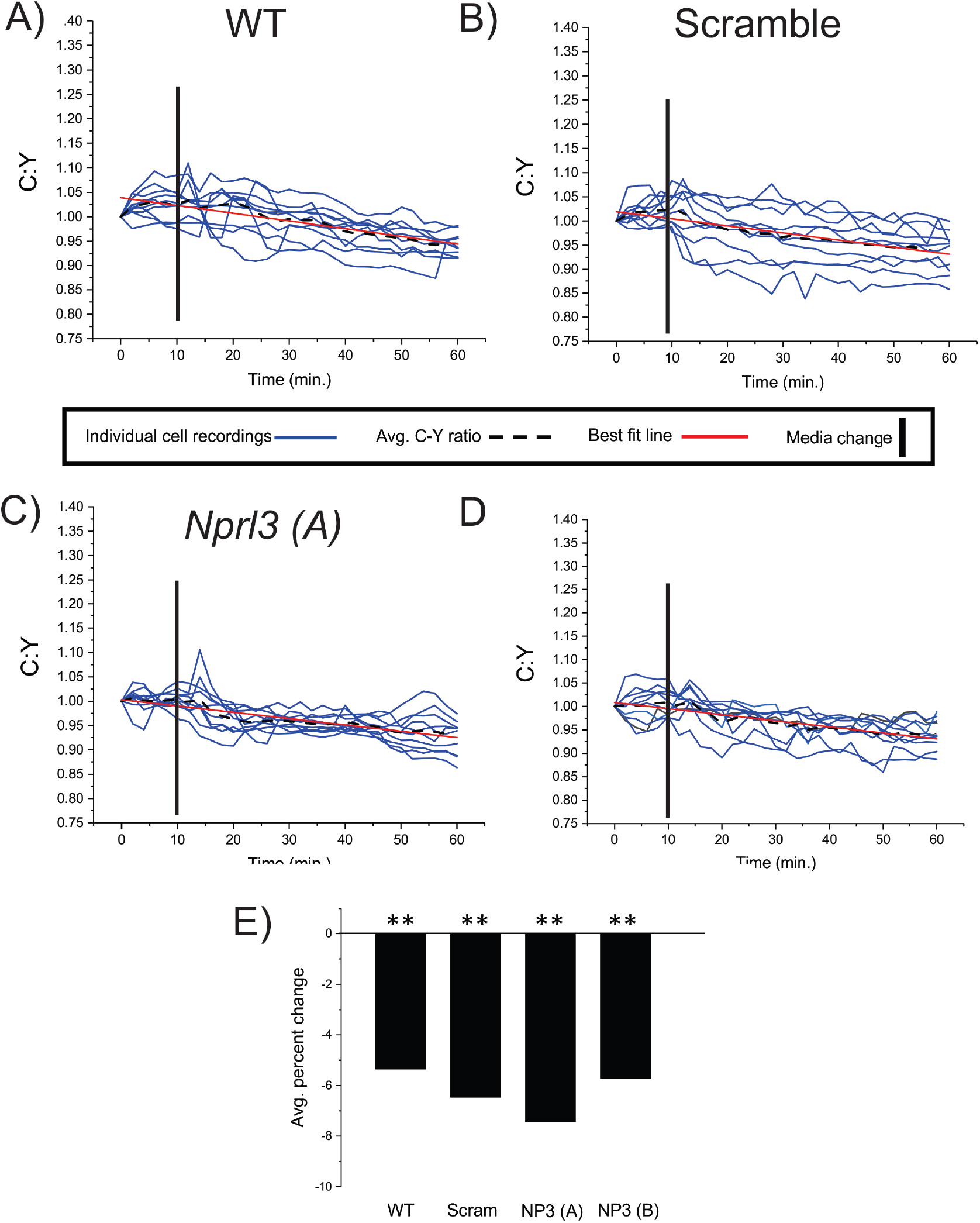
There is a decrease in 4E-BP1 phosphorylation at the lysosome during torin1 treatment of WT, scram, and *Nprl3* KO cells. C:Y was measured in *Nprl3* KO, WT and scramble N2aC lines transfected with TORCAR. Cells were incubated in complete media for 60 minutes containing 100 nM torin1 taking measurements at two-minute intervals. A −5% and −6.2% decrease in C:Y was observed in WT and scramble N2aC lines during the recording period (A,B,E; p<0.01). In *Nprl3* KO N2aC a −7.4% and −5.5% decrease was observed in *A* and *B* lines, respectively (C,D,E; p<0.01). Statistical significance was determined in comparison to corresponding baseline recordings in Supp. Fig. 3. ** = p<0.01, WT= wildtype, scram= scramble, NP3 (A) = *Nprl3* A, NP3 (B)= *Nprl3* (B).

**Sup. Table A:**
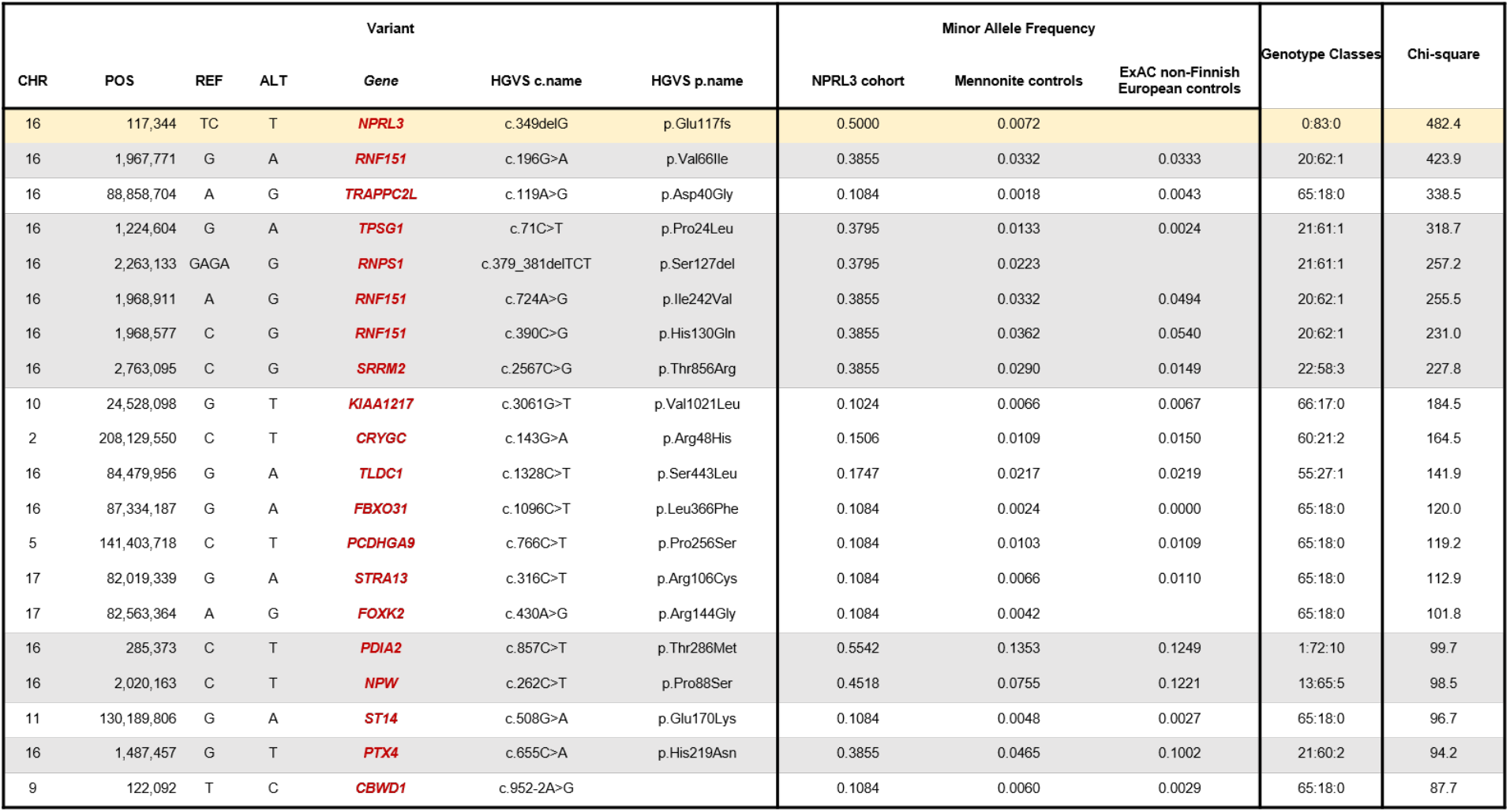
Genome-wide analysis of whole exome data for the entire *NPRL3* c.349delG cohort: For each variant identified in the *NPRL3* exome cohort (n=230,656), we calculated a chi-square statistic to assess allele frequency differences between heterozygotes and 835 Old Order Mennonite controls. This table provides information on the top 25 variants in this analysis. As expected, the *NPRL3* c.349delG variant (yellow shading) exhibited the greatest statistical allele frequency deviation. Nine additional variants (grey shading) from distal chromosome 16p13 also demonstrated highly significant chi-square values. The genotype classes are provided as follows - reference allele homozygote:heterozygote:alternate allele homozygote.

**Sup. Table B:**
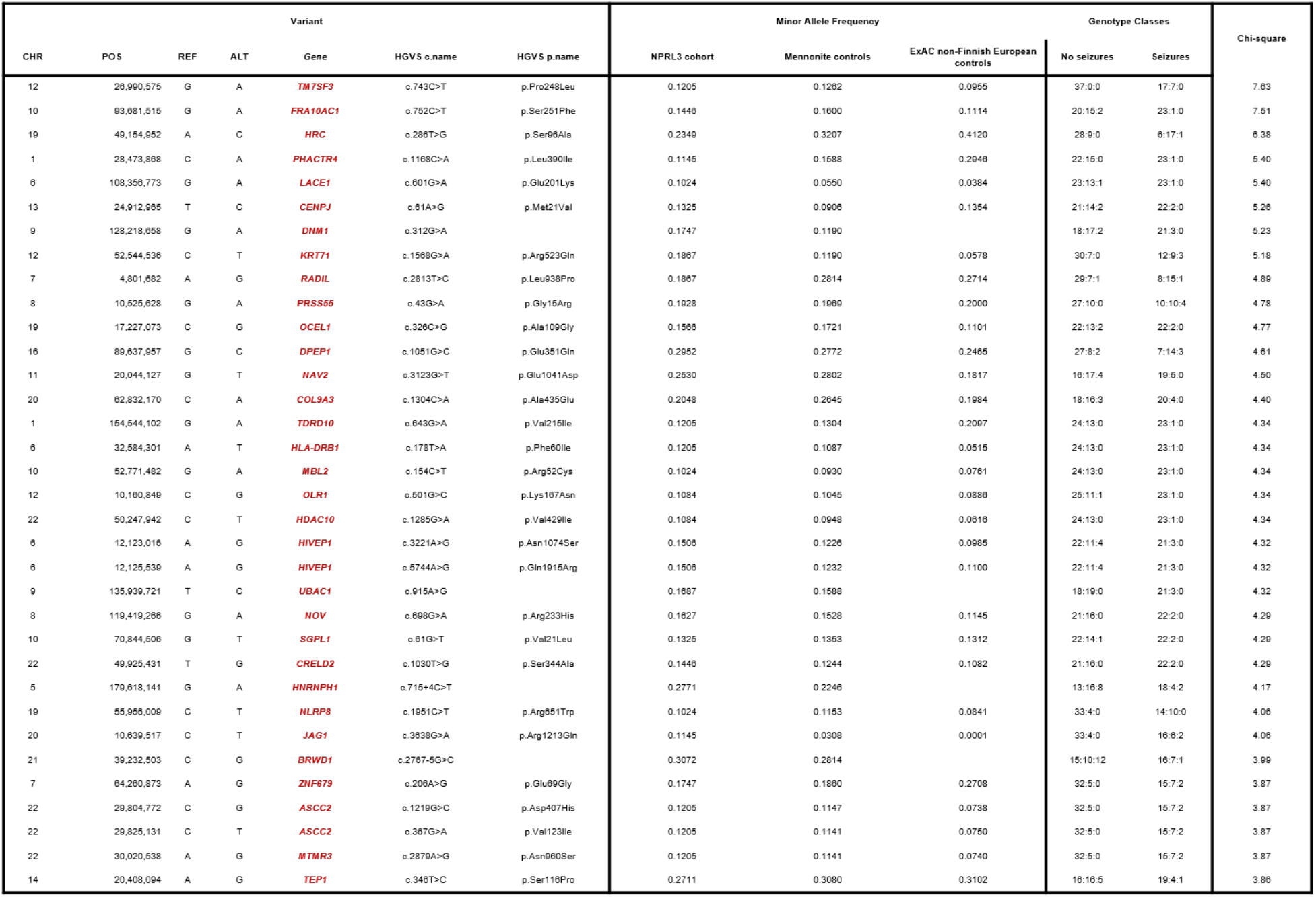
Genome-wide analysis of whole exome data in “seizures” versus “no seizures” *NPRL3* c.349delG heterozygotes: We calculated a chi-square statistic to assess allele frequency differences between *NPRL3* c.349delG heterozygotes with (n=24) and without seizures (n=37). This table provides information on all filtered variants with statistically significant (chi-square > 3.84) differences in allele frequency between the “seizure” and “no seizure” groups as described in the Results section. The genotype classes are provided as follows - reference allele homozygote:heterozygote:alternate allele homozygote.

**Sup. Table C:** Genome-wide analysis of whole exome data in “severe seizures” versus “no seizures” *NPRL3* c.349delG heterozygotes: We calculated a chi-square statistic to assess allele frequency differences between *NPRL3* c.349delG heterozygotes with severe seizures (n=10) and without seizures (n=37). This table provides information on all filtered variants with statistically significant (chi-square > 3.84) differences in allele frequency between the “severe seizure” and “no seizure” groups as described in the Results section. The genotype classes are provided as follows - reference allele homozygote:heterozygote:alternate allele homozygote.

| Variant |  |  |  |  |  |  | Minor Allele Frequency |  |  | Genotype Classes |  | Chi-square |
| --- | --- | --- | --- | --- | --- | --- | --- | --- | --- | --- | --- | --- |
| CHR | POS | REF | ALT | Gene | HGVSc c.name | HGVSp p.name | NPRL3 cohort | Mennonite controls | ExAC non-Finnish European controls | No seizures | Severe seizures |  |
| 12 | 26,990,575 | G | A | **TM7SF3** | c.743C>T | p.Pro248Leu | 0.1205 | 0.1262 | 0.0955 | 37:0:0 | 7:3:0 | 6.08 |
| 3 | 130,643,012 | G | A | **COL6A6** | c.5216G>A | p.Arg1739Gln | 0.1205 | 0.1878 | 0.1018 | 32:5:0 | 3:7:0 | 5.87 |
| 19 | 13,930,083 | G | A | **CC2D1A** | c.2716G>A | p.Ala906Thr | 0.1024 | 0.1298 | 0.1379 | 33:3:1 | 4:5:1 | 5.87 |
| 21 | 39,298,534 | G | C | **BRWD1** | c.247C>G | p.Gln83Glu | 0.3494 | 0.3877 | 0.4618 | 15:8:14 | 9:1:0 | 5.49 |
| 21 | 39,232,503 | C | G | **BRWD1** | c.2767-5G>C |  | 0.3072 | 0.2814 |  | 15:10:12 | 9:1:0 | 5.07 |
| 11 | 103,287,549 | C | T | **DYNC2H1** | c.11060C>T | p.Ala3687Val | 0.4096 | 0.3744 | 0.3561 | 8:18:11 | 8:2:0 | 4.68 |
| 5 | 179,618,141 | G | A | **HNRNPH1** | c.715+4C>T |  | 0.2771 | 0.2246 |  | 13:16:8 | 9:1:0 | 4.64 |
| 16 | 55,327,581 | C | T | **IRX6** | c.414-5C>T |  | 0.3855 | 0.3557 |  | 20:15:2 | 0:6:4 | 4.55 |
| 6 | 4,133,621 | C | T | **ECI2** | c.141G>A | p.Met47Ile | 0.1566 | 0.1812 | 0.2436 | 29:8:0 | 2:8:0 | 4.53 |
| 1 | 33,772,682 | T | C | **CSMD2** | c.1733A>G | p.His578Arg | 0.2470 | 0.2059 | 0.2552 | 23:5:9 | 10:0:0 | 4.50 |
| 3 | 186,671,770 | T | C | **HRG** | c.539T>C | p.Ile180Thr | 0.2349 | 0.1763 | 0.1226 | 20:12:5 | 10:0:0 | 4.26 |
| 17 | 68,269,226 | T | C | **SLC16A6** | c.1442A>G | p.His481Arg | 0.2711 | 0.2095 | 0.2435 | 16:12:9 | 9:1:0 | 4.22 |
| 3 | 81,648,979 | T | C | **GBE1** | c.568A>G |  | 0.1386 | 0.2295 | 0.3675 | 33:3:1 | 5:4:1 | 4.20 |
| 11 | 93,396,317 | G | C | **DEUP1** | c.1318G>C | p.Glu440Gln | 0.1145 | 0.1679 | 0.3531 | 32:5:0 | 7:0:3 | 4.20 |
| 11 | 93,396,318 | A | G | **DEUP1** | c.1319A>G | p.Glu440Gly | 0.1145 | 0.1679 | 0.3525 | 32:5:0 | 7:0:3 | 4.20 |
| 18 | 45,988,751 | T | C | **PSTPIP2** | c.964A>G | p.Asn322Asp | 0.1024 | 0.0996 | 0.1260 | 32:5:0 | 4:6:0 | 4.20 |
| 19 | 11,362,682 | C | G | **PLPPR2** | c.758C>G | p.Thr253Ser | 0.1446 | 0.1455 | 0.0720 | 32:5:0 | 5:4:1 | 4.20 |
| 8 | 17,960,044 | G | T | **PCM1** | c.2071G>T | p.Ala691Ser | 0.1988 | 0.1848 | 0.1558 | 25:10:2 | 3:3:4 | 4.18 |
| 1 | 46,033,854 | A | G | **MAST2** | c.3590A>G | p.Lys1197Arg | 0.1386 | 0.1051 | 0.0768 | 35:2:0 | 7:2:1 | 4.13 |
| 1 | 46,034,579 | G | A | **MAST2** | c.3910G>A | p.Val1304Met | 0.1386 | 0.1051 | 0.0788 | 35:2:0 | 7:2:1 | 4.13 |
| 12 | 52,544,536 | C | T | **KRT71** | c.1568G>A | p.Arg523Gln | 0.1867 | 0.1190 | 0.0578 | 30:7:0 | 5:3:2 | 3.94 |

**Supplemental Movie 1: Formation of cellular aggregates in complete media.** N2aC were plated at equal concentration in complete physiologic after passing through a cell strainer to eliminate any existing aggregates. Images were taken every 30 minutes for 48 hours and compiled into time-lapse videos in ImageJ. Multiple ROIs were chosen in each culture dish and representative time-lapse videos are shown. We observed large aggregates forming in both *Nprl3* KO lines, but only small aggregates were present in scramble or WT lines.

**Supplemental Movie 2: Formation of cellular aggregates is mTOR-dependent.** N2aC were plated at equal concentration in complete physiologic media containing 50 nM rapamycin after passing through a cell strainer to eliminate any existing aggregates. Images were taken every 30 minutes for 48 hours and compiled into time-lapse videos in ImageJ. Multiple sites were chosen in each culture dish and representative time-lapse videos are shown. Only rare aggregates formed in rapamycin treated ROIs during imaging.

**Supplemental Movie 3: Formation of cellular aggregates is prevented by torin1.** N2aC were plated at equal concentration in complete physiologic media containing 50 nM torin1 after passing through a cell strainer to eliminate any existing aggregates. Images were taken every 30 minutes for 48 hours and compiled into time-lapse videos in ImageJ. Multiple sites were chosen in each culture dish and representative time-lapse videos are shown. Only rare aggregates formed in torin1 treated ROIs during imaging.

